# Chromosome-Scale Genome Assembly of Australian Finger Lime and Resequencing Reveal the Hidden Diversity of Oceanian Citrus

**DOI:** 10.64898/2025.12.11.693794

**Authors:** Barbara Hufnagel, Monica Alves, Quentin Piet, Alexis Dereeper, Delphine Giraud, Lény Calvez, Caroline Belser, Karine Labadie, Simone Duprat, Corinne Cruaud, Benjamin Istace, Franck Curk, Fredson Dos Santos Menezes, Khadidiatou Diop, Maëva Miranda, Yann Froelicher, Malcolm Smith, Pablo Aleza, Stéphane Lebegin, Carole Martin, Fabienne Micheli, Pierre Mournet, Gaetan Droc, Leandro Peña, Nelson Wulff, Raphaël Morillon, Patrick Wincker, Jean-Marc Aury, Arnaud Lemainque, Patrick Ollitrault

## Abstract

Oceanian citrus species, including wild taxa native to Australia and Papua New Guinea, form a genetically distinct clade within the *Citrus* L. (1753) genus. These species remain largely underexplored, despite their adaptation to diverse environments and relevance for citrus improvement. To support their use in breeding and evolutionary studies, we generated a high-quality, Chromosome-scale genome assembly of an Australian finger lime accession (SRA 1002), a natural interspecific hybrid. The genome was assembled using long-read sequencing, optical mapping, and a high-density genetic map, resulting in nine pseudomolecules covering over 97% of the genome. Using this reference, we analyzed whole-genome resequencing data from 132 accessions representing the diversity of Asian and Oceanian citrus. Variant calling across the dataset produced a high-resolution catalogue of single nucleotide polymorphisms (SNPs) and derived database of SNP fully discriminant of 20 *Citrus* species (DSNPs), enabling detailed exploration of inter- and intra-specific diversity. The data reveal a strong population structure within the group with clear heterozygosity variation between ancestral species and admixed accessions, reflecting their complex evolutionary history and hybridization patterns. This study provides the first integrated genomic framework for Oceanian citrus diversity, offering essential tools for downstream applications in citrus breeding, conservation, and evolutionary genomics. The resources generated lay the groundwork for future association studies and the targeted introgression of beneficial traits into cultivated citrus.

## Introduction

Citrus is among the most important fruit crops worldwide, cultivated in more than 140 countries for its economic, nutritional, and cultural significance. The genus *Citrus*, as currently circumscribed (Ollitrault *et al*. 2020; Mabberley 2022), includes the cultivated gene pool resulting from a reticulate lineage of interrelated species that originated from a small number of ancestral taxa in Southeast Asia (Ollitrault, Curk and Krueger 2020). Extensive interspecific hybridization, coupled with long-standing clonal propagation and apomixis, has fixed elite agronomic traits in cultivated varieties but has also drastically eroded genetic diversity within modern breeding germplasm (Wu *et al*. 2018; Curk *et al*. 2022). This genetic narrowing limits the potential for developing resilient citrus cultivars, especially under emerging threats such as Huanglongbing (HLB), a devastating bacterial disease that threatens citrus production worldwide (Ghosh *et al*. 2022)

In contrast to the cultivated citrus gene pool, the Oceanian citrus species, comprising several taxa formerly classified under *Microcitrus Swin*gle (1915)*, Eremocitrus* Swingle (1914)*, Clymenia* Swingle *(1939)* and *Oxanthera* Montrouz (1860) (Swingle and Reece 1967), represent an underexplored reservoir of genetic and phenotypic diversity. These species now included in the *Citrus* genus (Ollitrault et al. 2020; Mabberley 2022), native to Australia, Papua New Guinea and New Caledonia, occupy diverse ecological niches ranging from the arid interior (*C. glauca* [Lindl.] Burkill [1932])) to the tropical rainforests of Queensland (*C. australasica* F. Muell., *C. australis* (Mudie) Planch (1858)., *C. inodora* F.M. Baileay (1889) (Nakandala *et al*. 2024a). They form a monophyletic group distinct from the Asian citrus lineages, as demonstrated by nuclear phylogenomic analyses (Nakandala *et al*. 2023b). Oceanian citrus species exhibit valuable agronomic traits including drought and cold tolerance, short juvenility, and most notably, resistance or tolerance to HLB (Ramadugu *et al*. 2016; Alves *et al*. 2021).

Recent studies have underscored the promise of these species in citrus improvement programs. Hybrids between Oceanian and Asian citrus have already been developed and evaluated, revealing the potential to introgress resistance traits into elite germplasm (Mahmoud and Dutt 2025) and to develop new rootstocks (Smith *et al*. 2024). Nevertheless, a comprehensive understanding of the genomic architecture and evolutionary dynamics of these wild species has been limited by the lack of high-quality, chromosome-scale reference genomes. Only in the past two years have phased genome assemblies been reported for several Australian taxa, including *C. australasica*, *C. australis*, *C. glauca*, and *C. inodora*, revealing structural variations, unique gene families, and lineage-specific expansions related to pathogen defense and stress adaptation (Nakandala *et al*. 2023b; Singh *et al*. 2024).

Among these species, *C. australasica* (Australian finger lime) has gained particular attention for its horticultural and culinary value, as well as its broad phenotypic variability in fruit morphology, color, and flavor (Delort & Yuan 2018). Its ease of hybridization with other *Citrus* species has made it a key parent in breeding programs aimed at improving fruit quality and disease resistance. However, preliminary GBS studies (our unpublished data under PreHLB, H2020 project) have shown that several Australian finger lime accessions maintained in French, Spanish, and Brazilian germplasm banks were introgressed with *C. australis*, forming complex interspecific mosaics. Understanding the genomic consequences of such introgression events and the distribution of structural variants across Oceanian lineages remains an open question.

Here, we present a chromosome-scale, high-contiguity genome assembly of Australian finger lime accession SRA 1002 (AFL SRA 1002), a suspected interspecific backcross hybrid *C. australasica* × (*C. australis* × *C. australasica*), maintained at the Citrus Biological Resource Center (INRAE-CIRAD) in Corsica (Luro *et al*. 2017). Using a combination of long-read Nanopore sequencing, PCR-free Illumina reads, Bionano optical maps, and a high-density interspecific genetic map, we anchored over 97% of the genome into nine pseudomolecules. We further analysed whole genome resequencing data of 132 accessions (including 54 new resequencing) representing the diversity of Asian and Oceanian *Citrus* to characterize population structure, detect interspecific introgressions, and identify diagnostic SNPs across clades.

This work provides the first integrated genomic framework for the study of Oceanian *Citrus* diversity. It establishes foundational resources for evolutionary analyses, comparative genomics, and breeding applications aimed at exploiting wild diversity for improving stress tolerance and disease resistance in cultivated Citrus.

## Results

### Australian Finger Lime sequencing and assembly

The reference genome of AFL SRA 1002 was assembled using a hybrid strategy that combined Oxford Nanopore long reads, PCR-free Illumina short reads, and Bionano optical maps. The initial assembly comprised 403 scaffolds totaling 450.8 Mb, with high contiguity (N50 = 28.4 Mb; N80 = 5.4 Mb) (Supplementary Table 1, Supplementary Note 1). K-mer analysis using GenomeScope 2.0 estimated a genome size of ∼338 Mb and a high heterozygosity rate, consistent with the hybrid origin of this accession (Supplementary Figure 1A).

The admixture analysis based on DSNPs (see “Diagnostic SNPs reveal clade differentiation and hybrid ancestry”) from WGS data of 132 accessions confirmed that AFL SRA 1002 is not a pure *C. australasica* but an interspecific hybrid, showing alternating homozygous *C. australasica* and heterozygous *C. australis* / *C. australasica* regions (Figure 1A). These findings support a backcross origin: *C. australasica* × (*C. australasica* × *C. australis*), consistent with the presence of *C. australasica* cytoplasm.

**Figure 1:**
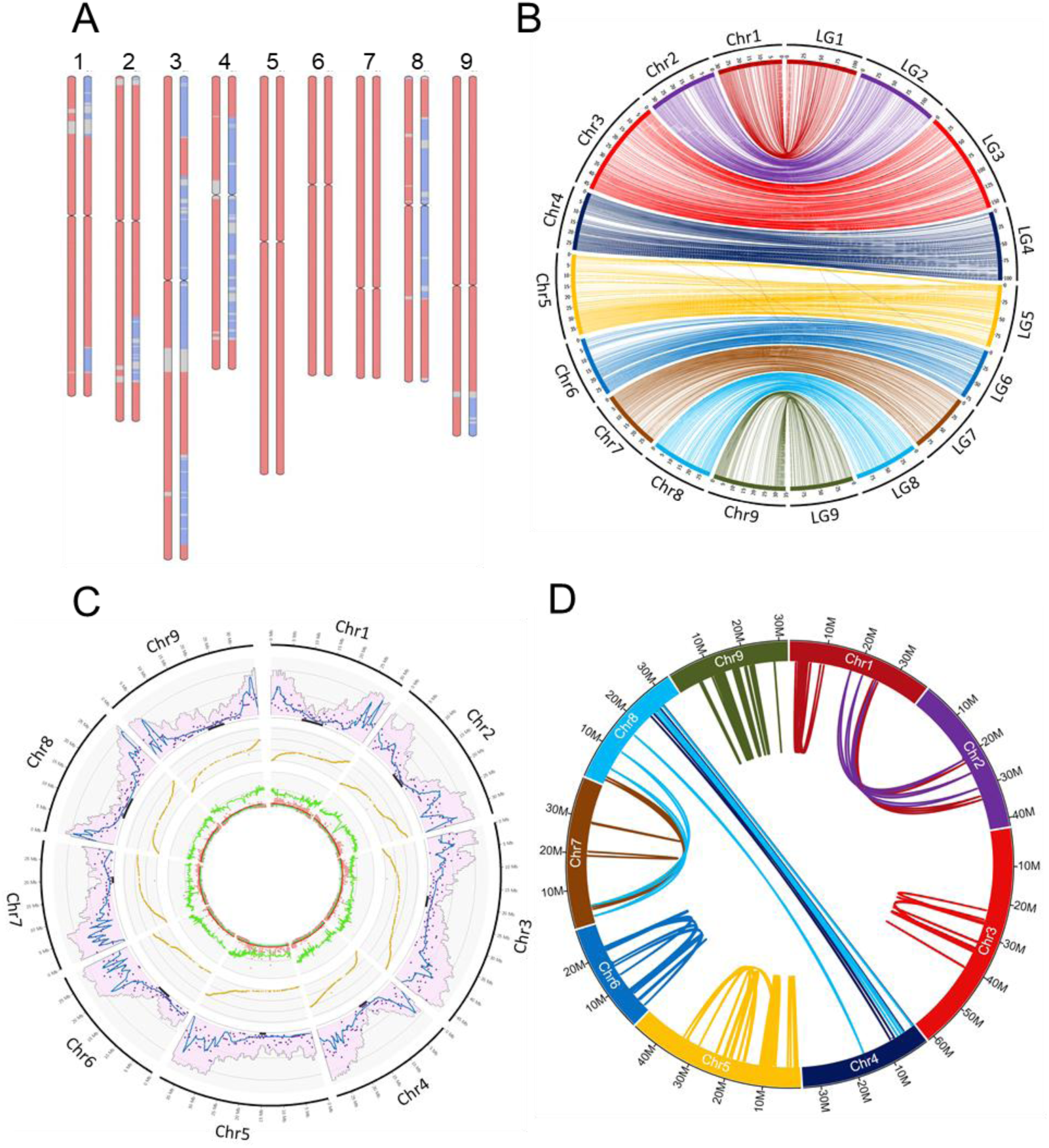
Overview of genome structure and quality metrics in AFL SRA 1002. **A)** Mosaic structure of AFL SRA 1002, which revealed a backcross origin: *C. australasica* × (*C. australasica* × *C. australis*); pink: *C. australasica*; light blue: *C. australis*; grey: undetermined. **B)** Anchoring of the AFL SRA 1002 assembly to the *C. australis* x *C. inodora* genetic map. Each ribbon connects a genomic position from the pseudochromosome assembly (Chr) to its corresponding position in one of the nine linkage groups (LG). The strong one-to-one correspondence and limited cross-linking support the structural accuracy of the assembly. **C)** Synthesis of parameter distribution along the genome. External layers: gene density (pink histogram), marker density (purple dots), and recombination landscape (blue line). Inner layers: Marey map (Y axis: genetic position) and segregation distortion analysis. Green line: *C. australis* haplotype contribution; brown dots: chi-square test q-values (based on expected 0.5/0.5 segregation); red line: significance threshold at q = 0.05. D) Intragenomic segmental duplications. Circular representation of syntenic blocks >7.6 kb identified by SynMap. Lines connect homologous segments between and within chromosomes.

Dotplot comparisons against the *C. reticulata* CITRE pseudochromosome assembly (Droc *et al*. 2024) revealed scaffold duplications, particularly in heterozygous interspecific regions (Supplementary Figure 1B). To correct for these redundancies, a curated assembly was generated by removing the duplicated scaffolds and reordering the remaining ones based on synteny and collinearity with the *C. reticulata* genome, in line with the conserved macrosynteny across Asian and Oceanian citrus species (Ollitrault *et al*. 2024).

The resulting pseudomolecule assembly comprised 13 scaffolds spanning 300.85 Mb, representing 96.86% of the total genome length. These were anchored into nine pseudomolecules. Unanchored scaffolds were concatenated with 100 "N" spacers to form a single unplaced scaffold ("Chr Un"), totaling 9.77 Mb. A first genetic map of a *C. australis × C. inodora* hybrid was produced using the *C. clementina* V1.0 reference genome from GBS data of a progeny of 171 ‘Fortune’ mandarin (*C. reticulata*) × (*C. australis × C. inodora*) hybrids (Ollitrault *et al*. 2024). We performed a new variant calling and genetic mapping from the same data using the new AFL assembly in pseudochromosome as template. 2,622 diallelic SNPs markers were successfully mapped in nine linkage groups encompassing 943.61 cM (Supplementary Table 2).

The genome assembly and genetic map were highly syntenic and collinear (Figure 1B) with only 9 markers (0.34%) discordant for synteny and a Spearman collinearity coefficient of 0.9997 ± 0.0001. These results validate the structural quality of the AFL genome assembly. The recombination landscape study revealed large genomic regions with very low recombination levels corresponding to low gene density areas (Figure 1C). The concordance between these two parameters was used to estimate the location of the centromeres (black lines on the basis of external figures of Figure 1C). Genomic regions with significant segregation distortion were observed in Chr 1, 3 and 5. In all cases they corresponded to areas with a deficit for transmission of *C. australis* haplotype.

The global synteny and collinearity of the AFL assembly and other *Citrus* species was checked by the anchoring of the consensus genetic map of the *Citrus* genus that include 10,756 markers (Ollitrault *et al*. 2024). It resulted in evidence for high synteny (96.72) and collinearity (Spearman coefficient = 0.9974 ± 0.0018); Supplementary Figure 1C).

### Genome assembly statistics and gene annotation

The final AFL genome assembly comprised nine pseudochromosomes totaling 300.8 Mb and an additional 9.7 Mb of unanchored scaffolds. Transcriptomic data from flowers, leaves, and fruit pulp (196.4 Gb total) were used for gene annotation. BUSCO completeness exceeded 91% in most tissues, except fruit pulp (76%) (Supplementary Table 3). Using the EuGene pipeline, 28,266 protein-coding genes were predicted (Table 1), distributed according to recombination-rich regions, consistent with other citrus genomes.

**Table 1.** Size and number of genes per pseudochromosome of the AFL genome.

| Pseudo-Chr | Size (bp) | Gene number |
| --- | --- | --- |
| 1 | 30,981,035 | 2,830 |
| 2 | 33,420,155 | 3,459 |
| 3 | 46,787,044 | 4,818 |
| 4 | 28,396,107 | 2,781 |
| 5 | 38,550,532 | 3,573 |
| 6 | 29,000,654 | 2,636 |
| 7 | 29,262,762 | 2,832 |
| 8 | 29,613,850 | 2,537 |
| 9 | 34,833,955 | 2,775 |
| Un | 9,772,391 | 25 |
| Total | 300,846,094 | 28,266 |

### Intragenomic segmental duplications in the AFL genome

The analysis of intragenomic duplications in the AFL SRA 1002 genome, using SynMap (Haug-Baltzell *et al*. 2017) with a minimum of five collinear genes per block, identified 1,379 duplicated genes grouped into 109 syntenic blocks. These blocks ranged in size from 37.8 kb to 1.69 Mb, with a mean length of 174.2 kb and a median of 124.5 kb. For visualization, only 131 blocks exceeding 7.6 kb were retained, corresponding to the upper 5% of duplication sizes (Figure 1D).

The resulting duplication landscape highlights a clear predominance of intrachromosomal duplications, particularly on Chr 9, which harbors the highest number of duplicated genes, and Chr 5, engages extensively in interchromosomal exchanges. Notably, a strong syntenic connection is observed between Chr 4 and Chr 8, suggesting potential ancient segmental translocations or duplications mediated by transposable elements. Beyond these structural patterns, a subset of duplicated genes encodes ABC transporters, oxidoreductases, and putative transcriptional regulators, reflecting functional diversification. Together, these findings support the role of segmental duplications in shaping the dynamic architecture of the AFL genome and in providing evolutionary raw material for subfunctionalization, neofunctionalization, or pseudogenization.

### Conservation of citrus repeat diversity in AFL SRA 1002, but also minor novelties

The annotation of repeated sequences revealed that AFL SRA 1002 chromosomes contained a large amount of TEs, representing 42.55% of the genome, with 31.09% of retrotransposons and 10.50% of DNA transposons (Figure 2A and Table 2). Among TEs identified in *C. australasica*, 99% has already been characterized in other citrus species (Giraud 2025). However, the *de novo* annotation allowed us to assemble 25 new consensus sequences of TEs found exclusively in AFL SRA 1002, covering 0.17% of the genome with 1,276 copies. These TEs are mainly LTR retrotransposons, LINEs and MITEs (9, 6 and 5 consensus, respectively; Supplementary Table 4). Interestingly, 363 and 449 truncated copies of two different LINEs were identified, suggesting ancient accumulation events in AFL SRA 1002 after its divergence from other citrus species. Further analyses will be conducted to determine the origin of these 25 TEs (new insertion events or derived from TEs already found in related citrus species) and their specific dynamics in AFL SRA 1002 and other Australian citrus species. The 25 consensus sequences were added to the citrus reference library of repeated sequences v2.0 (Giraud 2025).

**Figure 2:**
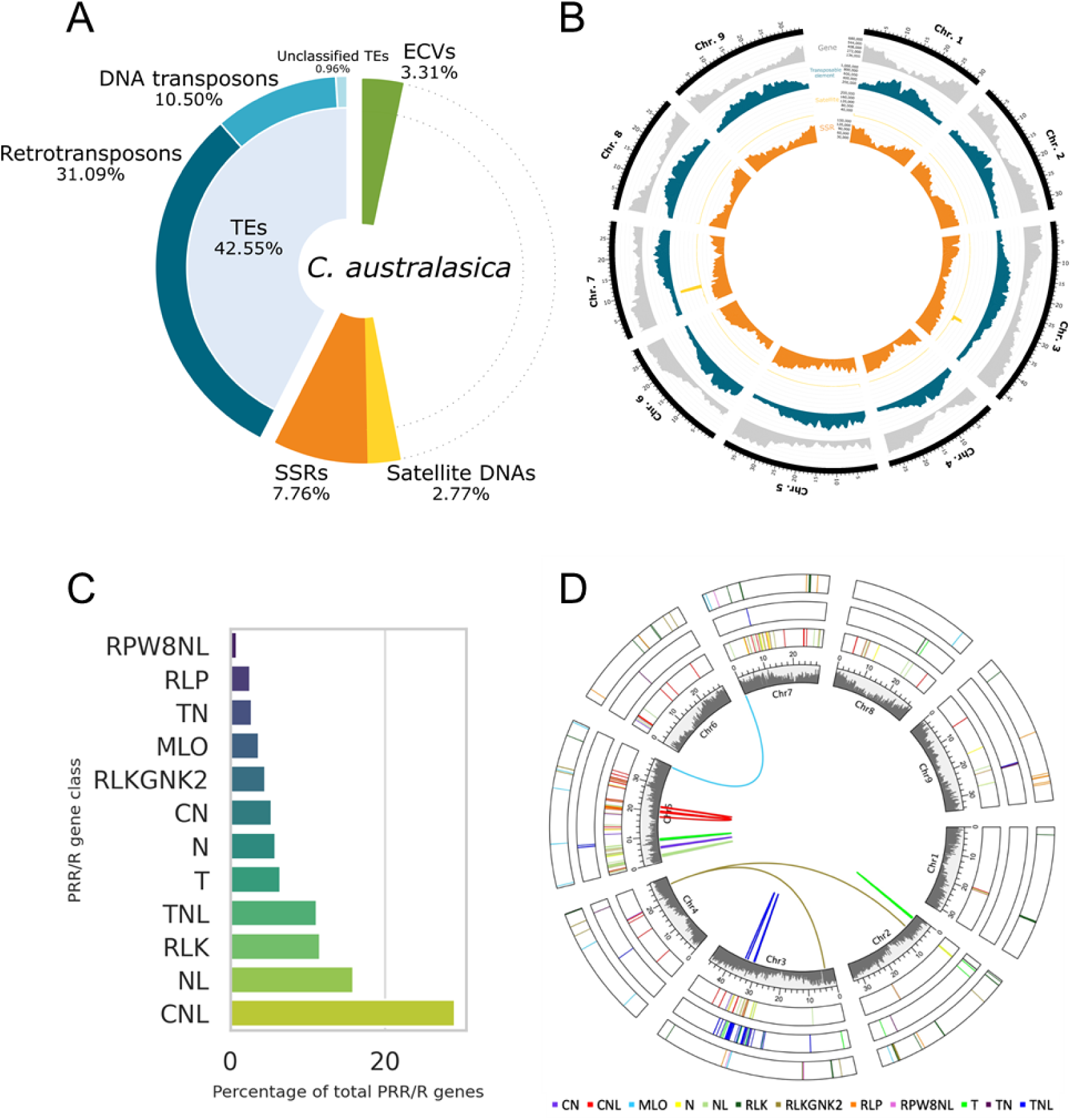
Annotation of repeated elements and immune receptor genes in the AFL SRA 1002 genome. **A**) Composition of repeated elements across the genome, including retrotransposons (LTRs and LINEs), DNA transposons, simple sequence repeats (SSRs), satellite DNAs, and endogenous caulimovirid elements (ECVs). **B)** Circos plot showing the chromosomal distribution of repeated elements in AFL SRA 1002. From outer to inner rings: (1) gene density; (2) transposable elements (TEs); (3) simple sequence repeats (SSRs); (4) satellite DNAs; (5) endogenous caulimovirid elements (ECVs). TEs are concentrated in centromeric and pericentromeric regions, while SSRs are enriched in gene-rich areas. **C)** Proportional representation of 12 well-defined classes of Pattern Recognition Receptors (PRRs) and resistance (R) genes annotated in the AFL SRA 1002 genome. **D)** Chromosomal localization and class distribution of PRR and R genes across the 12 well-established classes. Each colored arc represents a gene of a specific PRR/R class mapped onto the nine pseudochromosomes of AFL SRA 1002. Colored lines in the center of the figure: duplication events. Grey histograms represent gene density.

**Table 2:** Estimated quantities of repeated sequences in the AFL SRA 1002 genome assembly.

|  | Genome coverage (Mb) | Genome Coverage (%) |
| --- | --- | --- |
| **Retrotransposons (Class I elements)** |  |  |
| LTR Copia (RLC) | 45.38 | 14.61% |
| LTR Gypsy (RLG) | 42.97 | 13.83% |
| **LTR Total** | **88.35** | **28.44%** |
| LINE (RIX) | 7.66 | 2.47% |
| SINE (RSX) | 0.53 | 0.17% |
| Unclassified Class I | 0.04 | 0.01% |
| **DNA transposons (Class II elements)** |  |  |
| TIR elements (DTX) | 25.45 | 8.19% |
| MITE | 5.63 | 1.81% |
| Helitron (DHX) | 1.55 | 0.50% |
| Unclassified_TE | 2.98 | 0.96% |
| **TOTAL TE** | **132.18** | **42.55%** |
| ECVs (Endogenous caulimovirid elements) | 10.27 | 3.31% |
| **Satellite DNAs** | 8.60 | 2.77% |
| **SSR** | 24.10 | 7.76% |
| **TOTAL Repeated sequences** | **175.15** | **56.39%** |

As observed in other *Citrus* species, TEs were mainly concentrated in poor gene regions, including centromeres and pericentromeres and formed highly concentrated chromosomal arms in acrocentric chromosomes such as Chr 5 to Chr 7 (Figure 2B) also observed in Asian *Citrus* species (Droc *et al*. 2024). Clusters of satellite DNAs, which covered 2.77% of the genome, were also identified in TE-rich regions (Chr 3 and Chr 7) and in telomeres (Chr 1, Chr 5 and Chr 8). On the contrary, SSRs were more concentrated in gene-rich regions and made up 7.76% of the genome. Finally, the specific annotation allowed us to estimate that AFL SRA 1002 also contains a large number of endogenous caulimovirids elements (ECVs), representing 3.31% of its genome. The same diversity of ECVs has already been identified in other citrus species (Giraud *et al*. 2025), suggesting the vertical inheritance of its elements during citrus speciation.

### Diversity of Pattern Recognition Receptors (PRRs)

The annotation of the AFL SRA 1002 genome identified a total of 1,279 genes related to Pattern Recognition Receptors (PRRs) and resistance (R) genes. Among them, 463 genes (36.1%) were classified into 12 well-established categories, excluding the UNKNOWN class (Figure 2C; Table 3). Together, the CNL, TNL, and NL classes represent the canonical intracellular NLR gene repertoire, totaling 260 genes in AFL SRA 1002. The CNL class was the most represented, accounting for 29.1% of the defined genes (135), followed by NL (74 genes, 15.9%), RLK (54 genes, 11.6%) and TNL (52 genes, 11.2%).

**Table 3.**
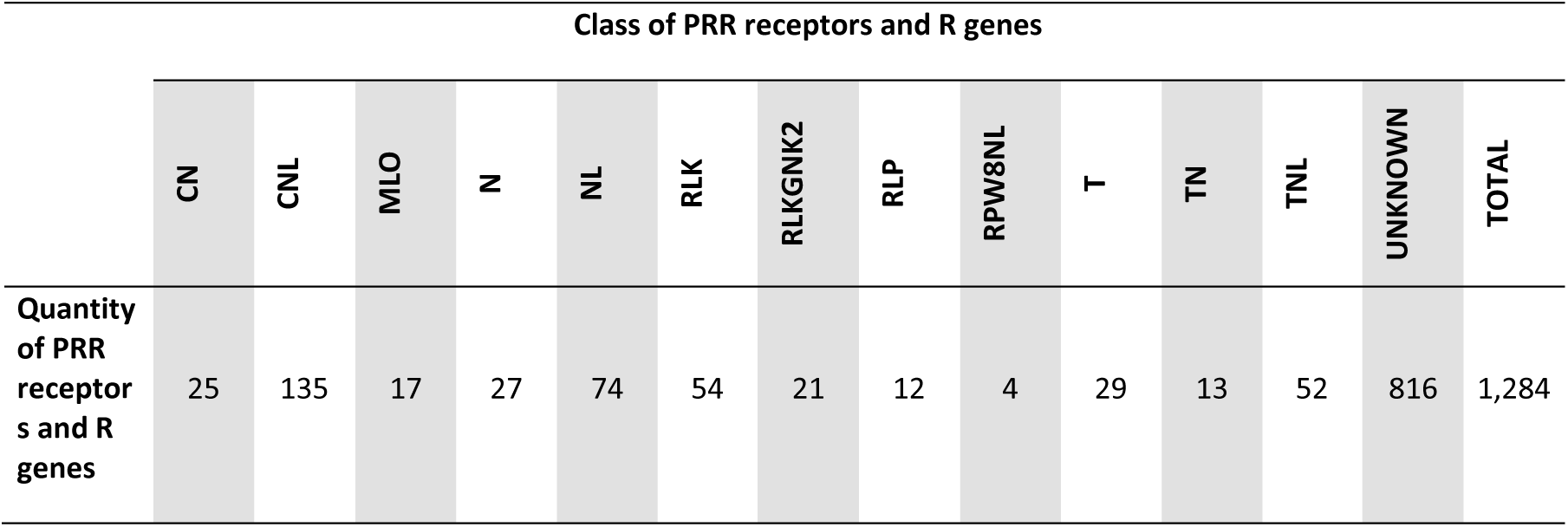
Quantity of PRR receptors and R genes present in AFL SRA 1002.

Other classes included N (27 genes, 5.8%), RLKGNK2 (21, 4.5%), CN (25, 5.4%), T (29, 6.5%), TN (13, 2.8%), MLO (17, 3.7%), RLP (12, 2.6%) and RPW8NL (4, 0.9%).

Chromosomal mapping revealed a non-uniform distribution of PRR/R genes, with the highest densities observed on Chr 5 (124 genes), 3 (116genes) and 7 (79 genes), and the lowest on Chr 4 (14), 1 (15) and 9 (19) (Supplementary Figure 2). These genes tended to cluster in gene-rich regions, particularly on Chr 2, 3 and 5, where the most represented classes – RLK, TNL, CNL and NL – were especially abundant (Figure 2D). Gene duplication analysis indicated several tandem duplication events, especially within the CNL and NL classes, each showing four duplication events (Figure 2D). Notably, the genes Ciaus5g18130 and Ciaus5g16660 (CNL), and Ciaus4g25280 (NL) were each involved in two independent duplications, highlighting possible hotspots of immune gene expansion in the AFL genome (Supplementary Table 5). By contrast, no duplication events were detected in the N, RLP, RPW8NL and TN classes. These findings reveal a structurally and evolutionarily dynamic PRR/R gene repertoire, which could underlie specific immune adaptations in *C. australasica* and *C. australis*.

### Genomic mosaicism and structural variation reveal untapped diversity in Oceanian citrus

Comparative structural analysis between the newly assembled AFL SRA 1002 genome and the published genomes of C. *australasica* (Singh *et al*. 2024) and *C. australis* (Nakandala *et al*. 2024b) revealed extensive conservation of syntenic blocks across all chromosomes, as well as multiple structural rearrangements such as inversions, translocations and duplications (Figure 3A). While the overall collinearity between AFL and both species was high, several large-scale discontinuities were observed, particularly in regions that failed to align or exhibited complex rearrangements, suggesting lineage-specific structural variation in AFL. These patterns highlight a mosaic genomic architecture in AFL, composed of large syntenic segments alternated with rearranged blocks, originating from both *C. australis* and *C. australasica* lineages. This structural configuration is consistent with an interspecific hybrid origin or a backcross scenario, as previously hypothesized from its nuclear mosaic signature and cytoplasmic inheritance.

**Figure 3:**
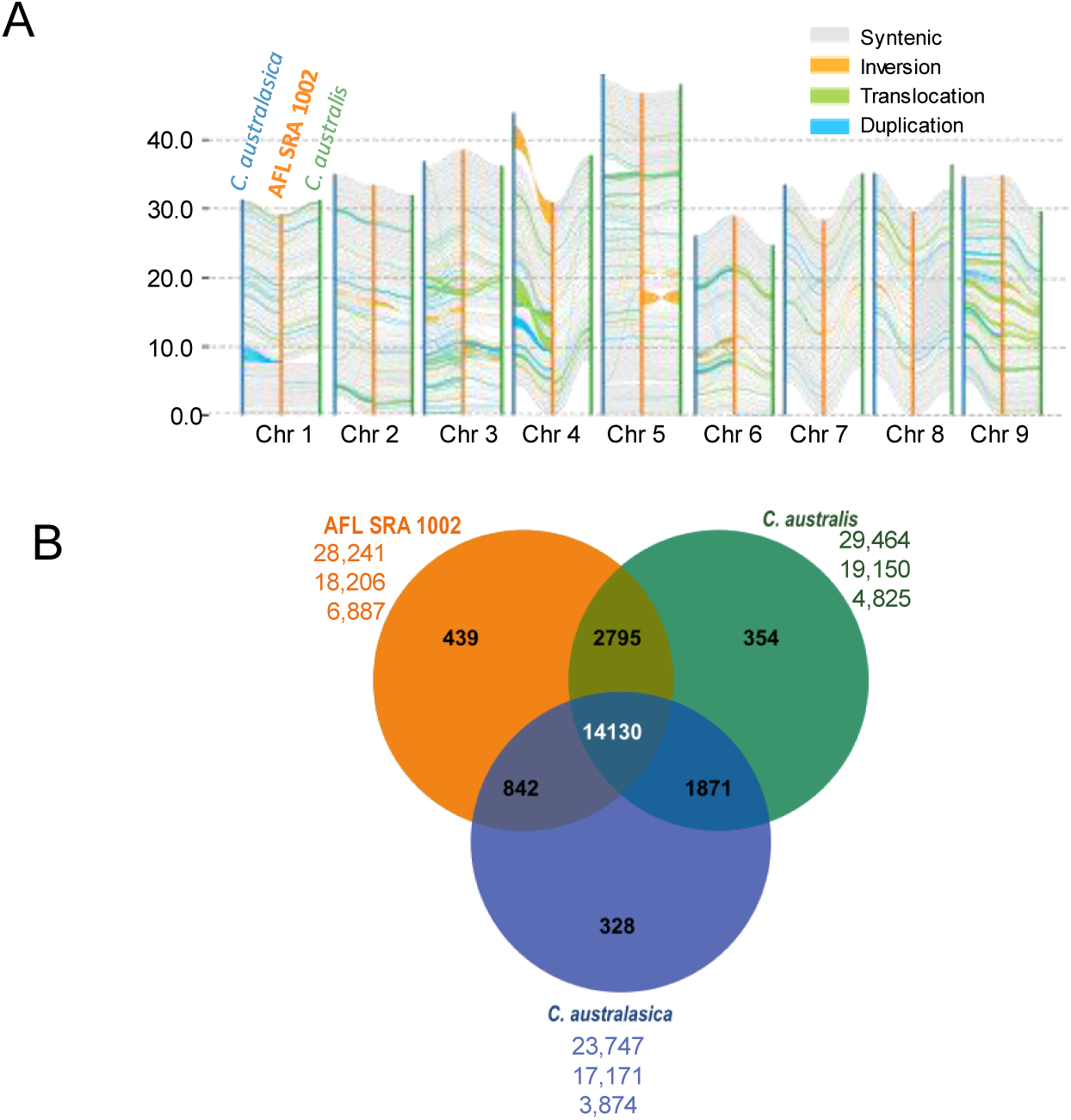
Structural genome comparison and gene clustering between AFL SRA 1002, *C. australasica*, and *C. australis*. **A)** Genome-wide structural comparison between AFL SRR 1002, *C. australasica* (primary haplotype; Singh et al., 2024), and *C. australis* (Nakalanda et al., 2023). Conserved syntenic blocks are shown in grey; non-aligned regions appear in white. Structural rearrangements (inversions, translocations, duplications) are color-coded by type. Analysis was performed using SyRI. B) OrthoMCL gene clustering among AFL SRR 1002, *C. australasica*, and *C. australis*. Venn diagram values represent the number of orthologous gene clusters. Below each species name, three values are shown: total number of genes, number clusters in each species, and number of unclustered genes.

Further comparisons with the two haplotype-resolved assemblies of *C*. *australasica* from Nakandala *et al*. (2024) (Supplementary Figure 3) revealed asymmetric synteny between AFL and each haplotype, reinforcing the hypothesis of complex hybrid ancestry. Despite the overall conservation, AFL displayed a distinctive pattern of rearrangements that were not shared with either haplotype alone, which may reflect recombination or selection processes following interspecific admixture. Gene clustering analysis using OrthoMCL (Figure 3B) supported these findings: Of the 28,241 predicted chromosomal genes in AFL, 21,354 were assigned to 18,206 orthologous clusters. Most clusters included genes from *C. australasica* and/or *C. australis*, and 439 clusters were specific to AFL. The remaining 6,877 genes did not cluster with any orthologs. These unclustered genes may correspond to lineage-specific sequences or highly diverged alleles not shared with the current *C. australasica* and *C. australis* references, or discrepancies arising from differences in genome assembly and gene annotation strategies. Their abundance, along with the observed structural variations, suggests a broader genetic diversity within Oceanian *Citrus* species than currently represented in available genome assemblies. These results indicate that additional diversity likely remains to be documented, particularly among under-sampled or unsequenced Oceanian species accessions.

### Genome-wide diversity and phylogenetic relationships

Variant calling was performed from WGS data of 132 citrus accessions (Supplementary Table 6) using the AFL SRA 1002 reference genome. A total of 40,698,669 SNPs and 6,585,258 small indels were identified (Table 4). The SNPs were distributed relatively evenly across the nine pseudochromosomes, with Chr 3 showing the highest SNP count (6.61 million) and Chr 6 the lowest (3.79 million). The SNP density along the genome was also relatively homogeneous, with however, lower density in the first half of chromosome 5 and the pericentromeric region of Chr 9 (Supplementary Figure 4).

**Table 4:** Total SNPs and indels per chromosome identified in 132 citrus accessions. Filtering for less than 15% of missing data).

| Chromosome | SNPs | Indels | Total Polymorphisms |
| --- | --- | --- | --- |
| Chr 1 | 4,452,695 | 704,811 | 5,157,506 |
| Chr 2 | 4,736,827 | 766,943 | 5,503,770 |
| Chr 3 | 6,612,355 | 1,085,099 | 7,697,454 |
| Chr 4 | 4,300,426 | 692,890 | 4,993,316 |
| Chr 5 | 4,529,864 | 742,567 | 5,272,431 |
| Chr 6 | 3,796,284 | 610,517 | 4,406,801 |
| Chr 7 | 4,048,772 | 671,818 | 4,720,590 |
| Chr 8 | 3,994,204 | 644,337 | 4,638,541 |
| Chr 9 | 4,209,553 | 664,571 | 4,874,124 |
| ChrUn | 17,689 | 1,705 | 19,394 |
| **Total** | 40,698,669 | 6,585,258 | 47,283,927 |

The analysis of heterozygosity along the genome reveals unimodal distribution with a peak under 0.005 for all pure representatives of ancestral Asian and Oceanian species (Supplementary Figure 5A and 5B). It is also the case for the two *C. wakonai* P.I. Forst. & M.W. Sm. (2010) and of the accessions “*C*. sp. Cape York,” (unknown), discarding the possibility that they derive from recent interspecific hybridization (Supplementary Figure 5C). Among the other accessions, some display a unimodal distribution with peak around 0.008 (mic-02fuD; supposed *C. australis × C. australasica* hybrid), 0.011 (oce-02fuD; PapCRC4111T; supposed *C. wintersii* Mabb. (1998) hybrids, Supplementary Figure 5D) and 0.017 (hEg-02fuD, hEg-orc2D and hEG-orfuD; supposed *C. glauca × C.* × *aurantium* L. (1753) hybrids, Supplementary Figure 5E) revealing their F1 interspecific status and an increasing divergence between parental species. The other accessions display multimodal heterozygosity distribution, suggesting complex admixture structures (Supplementary Figure 5F).

A filtered subset of 1,322,933 high-confidence diallelic SNPs covering the whole genome (with less than 5% missing data and ≥100 bp spacing) was used to build a nuclear phylogeny of the 103 accessions representatives of ancestral species. The tree revealed a differentiation supported by a bootstrap value of 100% between two Continental clusters with all Asian species (AsCl) one side and all Oceanian (OcCl) the other side. Sub branching within each Continental cluster lead us to identify 17 well-supported (bootstrap 100%) clades: 10 Oceanian and 7 Asian. (Figure 4, Supplementary Figure 6).

**Figure 4:**
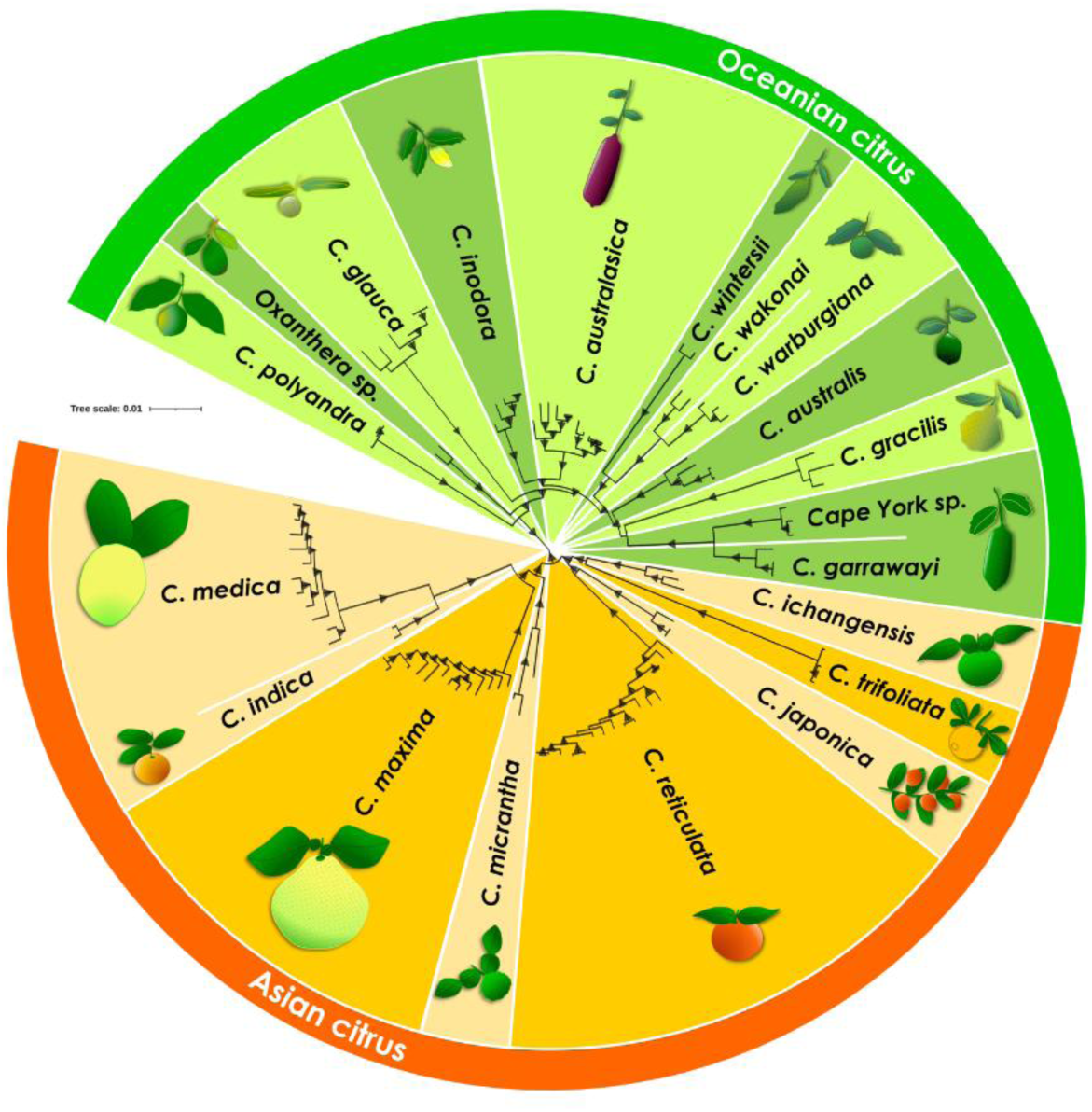
Nuclear phylogeny of 103 citrus accessions based on 1,322,933 genome-wide SNPs. The maximum likelihood tree identifies 17 well-supported clades, including 10 Oceanian lineages (green) and 7 Asian lineages (orange). A midpoint-rooted maximum likelihood tree was inferred using IQ-TREE v2.2.6 under the best-fit substitution model selected by ModelFinder Plus. Triangles indicate branch support (Boostrap >90; 1000 replicates), with size proportional to support values. The scale bar represents 0.01 substitutions per site.

*C. polyandra* (Ridel.) Burkill (1931) and *Oxanthera* Montrouz (1860) species that are both highly homozygous (Supplementary Figure 5) constitute two of the Oceanian clades and are the more basal of OcCl clade. *Citrus glauca* (desert lime, ADL), join then OcCl and occupies a distinct and isolated position within the tree with a high differentiation. All species of the former *Microcitrus* genus (Swingle and Reece, 1967) appears to be monophyletic. Within this “*Microcitrus*” cluster the three species from Papua New Guinea (*C. wintersii*, *C. warburgiana* F.M. Bailey (1901) and *C. wakonai* constitute a monophyletic group that join an Australian cluster constituted by *C. gracilis* Mabb. (1998) clade, *C. australis* clade and a former clade joining *C. garrawayi* F.M. Bailey (1900) and the three unclassified accessions from Cape York (designed as Unknown in this paper). Among the Papua New Guinean species, *C. wakonai* and *C. warburgiana* appear to be closely allied both at nuclear and chloroplast level (Supplementary Figure 7) and were included in a single main clade while *C. wintersii* constitute a second Papua New Guinea main clade. The Australian finger limes group (*C. australasica)* is supported as an independent lineage and a main clade of our analysis. It clusters with the last Oceanian main clade, *C. inodora,* with 100 % bootstrap value.

For Asian species seven main clades are distinguished with *C. medica* L. (1753) and *C. indica* Tanaka (1929) forming one of them at both nuclear and chloroplast levels. The rest of the organization is congruent with previously published phylogeny of Asian citrus performed from WGS data (Wu *et al*. 2018).

The chloroplast phylogeny, reconstructed using 3,695 SNPs (missing data > 5%) aligned to the *C.* × *sinensis* (L.) Osbeck (1765) plastid genome (Bausher et al. 2006), fully confirmed the 10 Oceanian and seven Asian main clades (Supplementary Figure 7).

However, the tree structure is different from the nuclear one as already described by Wu *et al*. (2018) and Nakandala *et al*. (2023a). It may be explained by potential reticulate evolution, incomplete lineage sorting, or hybridization/introgression.

Overall, the high-resolution phylogenetic reconstruction highlights a clear divergence between Oceanian and Asian citrus. It provides a robust framework for evolutionary and breeding studies and supports the clade structure observed through complementary organellar analyses.

### Diagnostic SNPs reveal clade differentiation and hybrid ancestry

To refine the resolution of diversity patterns, we identified diagnostic SNPs (DSNPs) for the main Oceanian and Asian clades inferred from nuclear and chloroplast phylogenies. Ten Oceanian clades were considered (*C. wakonai* + *C. warburgiana* grouped; *C. garrawayi* + Unknown Cape York sp. treated jointly) and seven Asian clades (*C. medica* + *C. indica* grouped).

Across the 17 clades, 4,390,992 DSNPs were identified, with 2,617,304 in Oceanian and 1,689,010 in Asian clades (Table 6). Within Oceania, the highest DSNP counts occurred in *C. polyandra* (546,168), *C. gracilis* (489,820), *C. wintersii* (439,052), and *C. glauca* (411,918), while *C. australasica* showed the fewest (20,067), likely reflecting higher intraspecific diversity. DSNPs were distributed along all nine chromosomes, concentrated in gene-rich regions (Supplementary Figs 8–9). Additionally, 84,678 DSNPs distinguished all Asian from all Oceanian accessions, providing a powerful genomic resource for tracing ancestry and lineage-specific traits (Table 5).

**Table 5:** DSNPs for the 17 main *Citrus* clades. Number of DSNPs identified per clade and chromosome.

|  | Chr 1 | Chr 2 | Chr 3 | Chr 4 | Chr 5 | Chr 6 | Chr 7 | Chr 8 | Chr 9 | Total |
| --- | --- | --- | --- | --- | --- | --- | --- | --- | --- | --- |
| ADL | 50 589 | 51 316 | 71 980 | 48 994 | 37 345 | 44 101 | 35 674 | 37 700 | 34 219 | 411 918 |
| AFL | 2 369 | 2 619 | 3 876 | 2 483 | 672 | 1 943 | 1 444 | 2 844 | 1 817 | 20 067 |
| ARL | 14 670 | 13 408 | 16 264 | 14 037 | 3 748 | 12 292 | 8 429 | 8 575 | 6 778 | 98 201 |
| CLY | 59 984 | 67 997 | 93 170 | 61 131 | 55 223 | 53 891 | 54 341 | 52 882 | 47 549 | 546 168 |
| GAR/Un | 22 401 | 21 506 | 29 856 | 20 850 | 14 220 | 19 541 | 13 264 | 17 744 | 12 860 | 172 242 |
| GRA | 64 634 | 73 831 | 72 527 | 58 331 | 55 168 | 45 163 | 45 655 | 40 234 | 34 277 | 489 820 |
| INO | 7 323 | 8 072 | 12 074 | 8 435 | 6 833 | 6 832 | 5 377 | 7 610 | 6 929 | 69 485 |
| OXA | 20 972 | 23 484 | 31 933 | 21 623 | 19 544 | 19 745 | 18 872 | 20 100 | 17 889 | 194 162 |
| WAR/W<br>AK | 20 943 | 23 205 | 29 034 | 21 601 | 14 350 | 19 460 | 15 614 | 15 965 | 16 017 | 176 189 |
| WIN | 49 491 | 58 555 | 69 793 | 56 237 | 38 151 | 41 755 | 42 947 | 41 034 | 41 089 | 439 052 |
| **Total**<br>**Oceanian** | **313 376** | **343 993** | **430 507** | **313 722** | **245 254** | **264 723** | **241 617** | **244 688** | **219 424** | **2 617 304** |
| ICH | 10 992 | 12 364 | 15 135 | 10 697 | 7 639 | 10 990 | 7 339 | 8 425 | 7 844 | 91 425 |
| JAP | 19 429 | 20 368 | 29 342 | 20 849 | 16 326 | 18 448 | 15 084 | 17 049 | 12 650 | 169 545 |
| MAX | 14 871 | 15 202 | 20 329 | 14 187 | 10 974 | 13 270 | 11 517 | 10 558 | 9 531 | 120 439 |
| MED/IN<br>D | 29 791 | 34 580 | 44 807 | 31 879 | 28 119 | 27 076 | 27 299 | 23 916 | 24 684 | 272 151 |
| MIC | 9 175 | 11 259 | 14 967 | 10 771 | 8 545 | 10 033 | 8 200 | 8 944 | 7 043 | 88 937 |
| RET | 20 354 | 21 060 | 29 189 | 19 135 | 15 915 | 17 991 | 13 922 | 15 829 | 13 776 | 167 171 |
| TRI | 87 199 | 100 508 | 132 142 | 88 793 | 75 223 | 78 312 | 72 359 | 75 099 | 69 707 | 779 342 |
| **Total**<br>**Asian** | **191 811** | **215 341** | **285 911** | **196 311** | **162 741** | **176 120** | **155 720** | **159 820** | **145 235** | **1 689 010** |
| Asian/<br>Oceanian | 10 116 | 10 765 | 14 942 | 10 481 | 8 512 | 8 076 | 7 405 | 8 065 | 6 316 | 84 678 |
| **Total** | **515 303** | **570 099** | **731 360** | **520 514** | **416 507** | **448 919** | **404 742** | **412 573** | **370 975** | **4 390 992** |

Species within shared main clades also displayed specific DSNPs: *C. warburgiana* + *C. wakonai* (93,807 and 85,026 DSNPs), *C. garrawayi* + Unknown Cape York sp. (65,420 and 100,055), and *C. medica* + *C. indica* (239,123 and 133,783). Their genome-wide distribution (Supplementary Table 7, Supplementary Fig. 10) supports the validity of these taxa, consistent with their low heterozygosity and the chloroplast phylogeny.

Using the AFL reference genome as a mapping backbone, DSNPs refined phylogenomic resolution and enabled detection of hybrid ancestry. Among Oceanian species, interspecific introgressions were detected only in *C. australasica*, where four accessions, including SRA 1002, displayed heterozygous *C. australasica* / *C. australis* regions (Supplementary Fig. 11), all sharing the *C. australasica* chlorotype (Supplementary Fig. 12). These *C. australasica* / *C. australis* admixtures correspond to *C. × virgata* (Mabb., 1998), as proposed by Mabberley (2022), encompassing accessions from Corsica and Brazil that exhibit complex hybrid structures consistent with selfing or backcrosses to *C. australis* (Fig. 5A).

**Figure 5:**
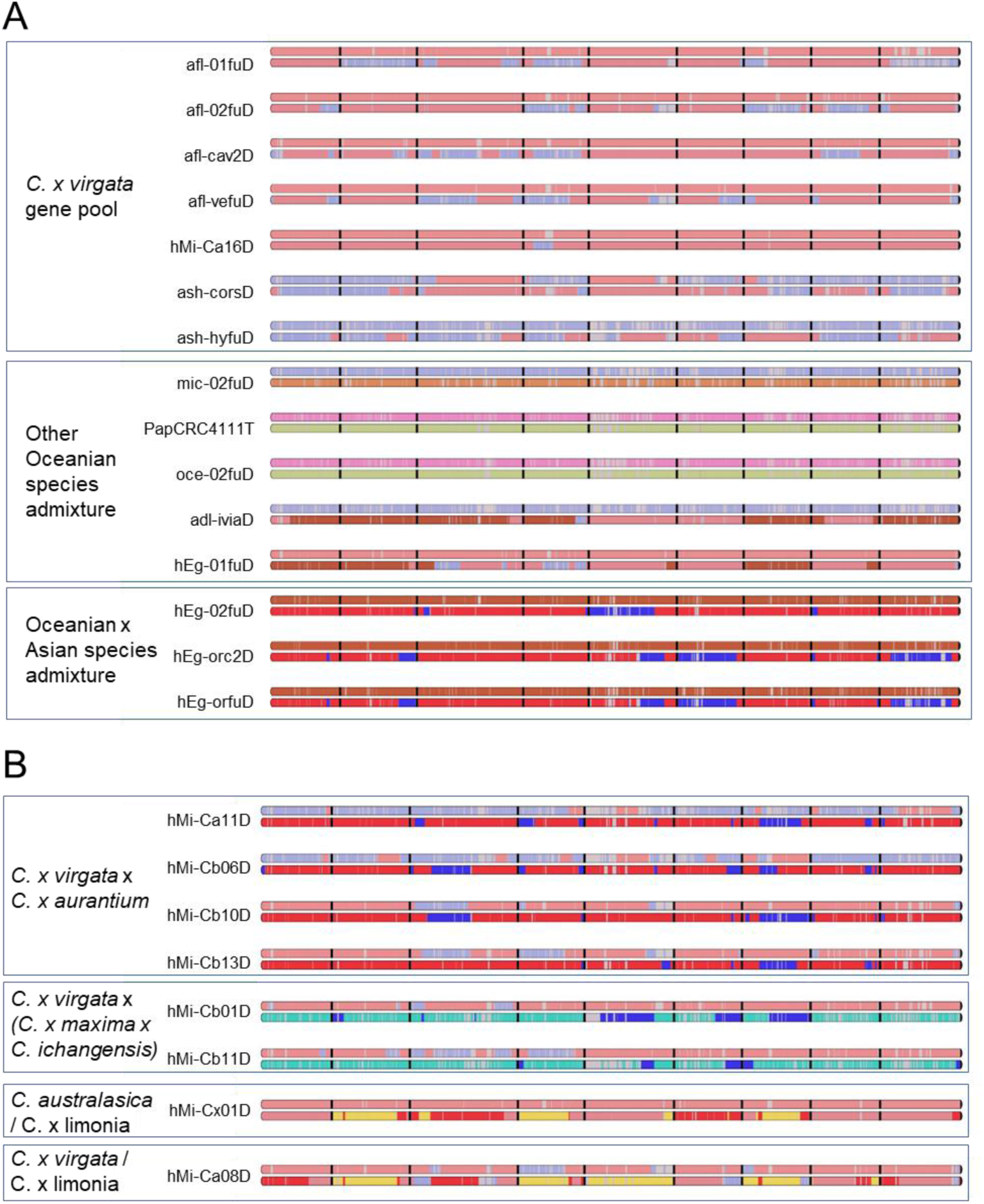
: Phylogenomic mosaic structure along the nine chromosome (separated by black dash) **A)** Admixed Oceanian citrus germplasm **B)** Aleatory admixed Oceanian/Asian hybrids of the CIRAD/INRAE breeding program. Color code: salmon: *C. australasica;* light blue: *C. australis,* light brown: *C. inodora*; green: *C. warburgiana*; pink: *C. wintersii*; dark brown: *C. glauca;* red: *C. reticulata*; blue: *C. maxima;* cyan: *C. ichangensis;* yellow: *C. medica*.

Other admixed genotypes included an F₁ *C. australis × C. inodora* (mic-02FuD), two F₁ *C. wintersii × C. warburgiana* (PapCRC411T, oce-02FuD), and complex mosaics combining *C. australasica*, *C. australis*, and *C. glauca* (adl-iviaD, hEg-01FuD), each with chlorotypes matching the maternal species (Supplementary Fig. 12). Eremorange hybrids (*C. glauca × C. × aurantium*) showed expected genomic mosaics of *C. glauca* and *C. reticulata* / *C. maxima*, with identical structures across Corsican and Brazilian accessions, confirming their identity (Supplementary Fig. 12). Asian admixed species exhibited mosaic genomic architectures consistent with previous analyses using Asian references (Droc et al., 2024) (Supplementary Fig. 13).

Further analyses of greenhouse-derived hybrids from random Oceanian × Asian pollinations revealed various generations of admixture (Fig. 5B). Four hybrids appear as F₁ between *C. × virgata* and *C. × aurantium* (all *C. reticulata* chlorotype), two as F₁ combinations of Oceanian and Asian genomes (*C. australasica*, *C. australis*, *C. ichangensis*, *C. maxima*; *C. australasica* chlorotype), and two advanced-generation mosaics likely involving *C. × limonia* (Asian side) and *C. australasica* or *C. × virgata* (Oceanian side), both with *C. reticulata* chlorotype (Supplementary Fig. 12).

Altogether, these results demonstrate that DSNPs not only delineate clades with high accuracy but also allow precise reconstruction of hybridization and introgression patterns, supporting germplasm authentication and guiding breeding strategies (Fig. 5A–B).

### Gene loss events and their functional implications

We performed a gene presence/absence variation (PAV) analysis using whole-genome sequencing (WGS) data mapped to the AFL SRA 1002 reference genome. After removing low-coverage accessions and those sequenced with Tell-Seq (to avoid bias), we retained 81 representative ancestral accessions for analysis. Using SGSGeneLoss with a minimum coverage threshold of 5× and a loss cutoff of 90%, we identified 123 genes absent in all accessions, including the reference genotype, likely reflecting assembly artifacts. These were excluded, and 28,143 genes were retained for downstream analysis. The factorial analysis based on gene loss profiles revealed a clear structuring of citrus diversity (Figure 6A). Asian citrus species clustered tightly together along the first axis, while *C. australasica* formed a distinct and homogeneous group. A Neighbor-Joining tree based on PAV data (Supplementary Figure 14) supported these observations, showing consistent specific clade structure for Asian species and a distinct *C. australasica* cluster.

**Figure 6.**
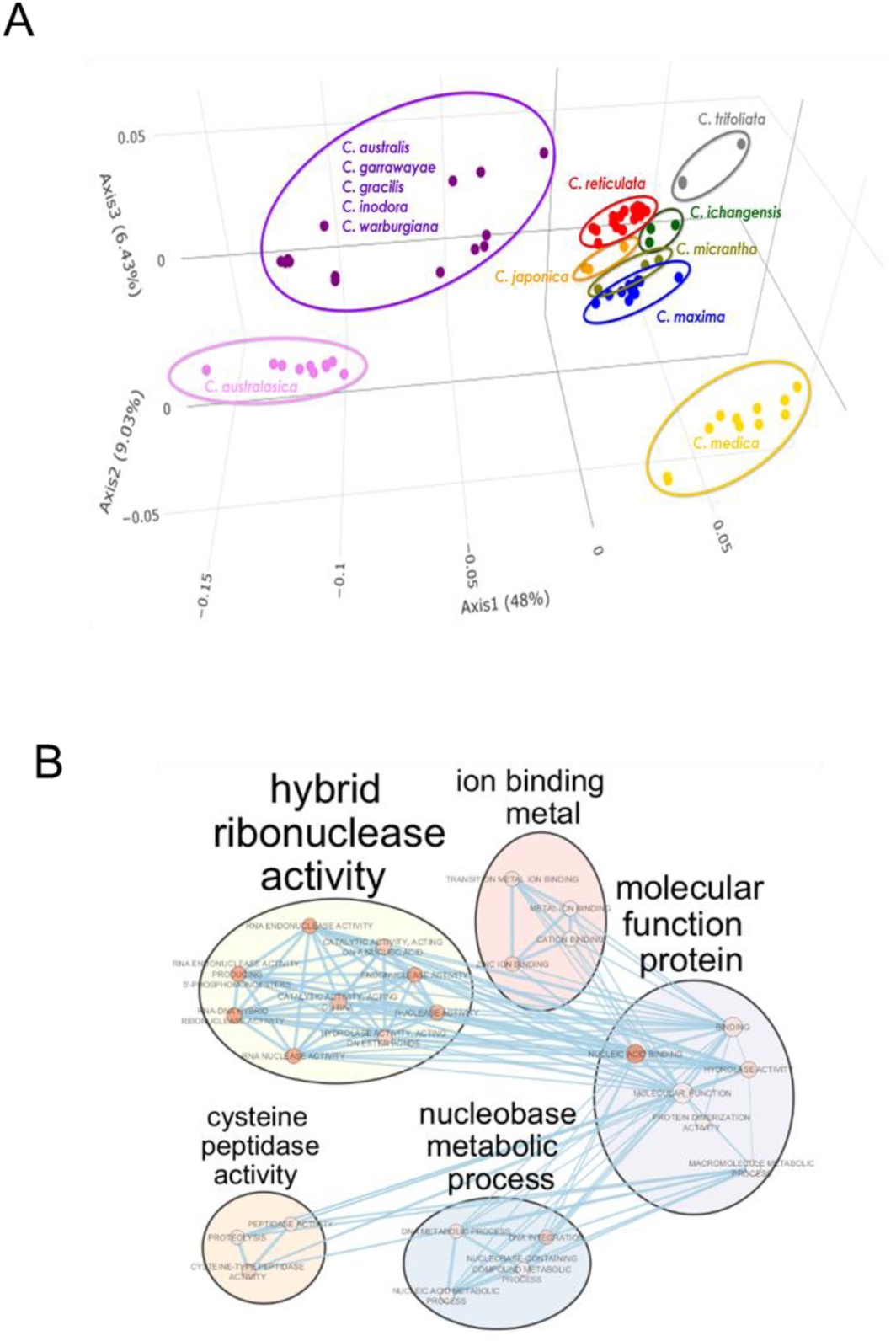
: Gene loss structuration and functional interpretation across Citrus diversity. **A)** Factorial analysis of 81 representative citrus accessions based on presence/absence variation (PAV) of genes mapped to the *AFL SRA 1002* reference genome. Asian Citrus species (e.g., *C. reticulata*, *C. maxima*, *C. medica*, etc.) cluster tightly along the first axis, while *C. australasica* forms a clearly distinct clade. Other Oceanian species (e.g., *C. australis*, *C. inodora*, *C. warburgiana*) show broader dispersion**. B)** GO enrichment network of the 1,090 genes absent in Asian Citrus species.

Among the 28,143 retained genes, 1,090 were completely absent in all Asian accessions. Of these, 127 were present in at least 95% of Australian and Papua–New Guinean accessions, and 52 were exclusive to *C. australasica*, suggesting potential species-specific adaptations. The flow of these filtered gene sets is summarized in a Sankey diagram (Figure 6B).

GO enrichment of the 1,090 genes absent in Asian citrus (Figure 6B) revealed an overrepresentation of functions related to nucleic acid metabolism, including RNA endonuclease and ribonuclease activities, as well as nucleus metabolic processes. Additional clusters highlighted metal ion binding and cysteine peptidase activity, which may be linked to oxidative stress regulation and protein turnover. These enriched functions suggest that genes, lost in Asian species but retained in Oceanian lineages, may contribute to broader metabolic plasticity, stress resilience, and genome stability.

To better characterize lineage-specific adaptations, we performed GO enrichment analysis on two subsets of genes absent in Asian citrus: the 127 genes conserved in ≥95% of Australian and Papua–New Guinean accessions, and the 52 genes exclusive to *C. australasica*. The 127-gene set showed significant enrichment in cation binding, DNA integration, and ribonuclease activity, supporting roles in genome stability and stress adaptation (Supplementary Figure 15A). In contrast, the 52 genes specific to *C. australasica* were enriched for UDP-glycosylation, nucleic acid catalytic activity, and RNA/DNA nuclease functions (Supplementary Figure 15B), consistent with specialized metabolic and defense-related processes potentially contributing to the species’ unique ecological resilience. Together, these results indicate that gene absence in Asian citrus may reflect both lineage-specific streamlining and the divergence of defense- and metabolism-related functions, while the retained genes in *C. australasica* and Oceanian relatives may underlie distinctive adaptive strategies.

## Discussion

In this study, we report a high-quality chromosome-scale genome assembly of the AFL SRA 1002, a backcross hybrid *C. australasica* x (*C. australasica* x *C. australis*) *with C. australasica* cytoplasm. The assembly integrates long-read Nanopore sequencing, PCR-free Illumina reads, optical mapping, and a high-density interspecific genetic map, resulting in nine pseudomolecules spanning over 97% of the estimated genome. The AFL SRA 1002 assembly shows very high synteny and collinearity with the *C. australis × C. inodora* genetic map and the consensus *Citrus* map (Ollitrault et al. 2024), confirming strong structural conservation across the genus. It reveals a high degree of synteny and collinearity of our assembly with other citrus genomes, such as *C. reticulata* (Droc *et al*. 2024). Comparative analyses with recently published genomes of *C. australis* (Nakandala *et al*. 2023b), and other two *C. australasica* (Nakandala *et al*. 2024b; Singh *et al*. 2024) accessions also confirm strong synteny across Oceanian citrus despite their ecological divergence.

Compared to previously published genomes of pure *C. australasica* and *C. australis* accessions (Nakandala *et al*. 2023b, 2024b; Singh *et al*. 2024), the SRA 1002 assembly represents a valuable hybrid genome that may reflects the recombined genetic mosaic present in breeding germplasm. The AFL SRA 1002 genome is therefore well suited for population-level analyses, QTL mapping, and candidate gene identification in interspecific hybrids aimed at resistance introgression.

Segregation distortion in the genetic map, particularly the counter-selection of *C. australis*-derived regions on Chr 1, 3, and 5, indicates postzygotic selection or gametic incompatibility, which could limit the recovery of certain hybrid genotypes. Understanding these transmission biases will be useful for guiding controlled hybridizations among Oceanian species.

The annotation revealed a TE landscape (42.55% of the genome) broadly consistent with other citrus genomes, but with distinct lineage-specific activity. Twenty-five new TE consensus sequences were found, mainly LTRs and LINEs, highlighting active or ancient transpositional bursts in Oceanian lineages. Future comparative annotation of TEs and endogenous caulimovirid elements (ECVs) across Australian species will help assess their contribution to genome evolution and gene regulation. The immune receptor repertoire (1 284 PRR and R genes) shows both conservation and expansion of major resistance classes, with CNLs, NLs, and TNLs predominating. The total number of NLRs (261) is lower than in *C. australasica*, *C. glauca*, and *C. inodora* (Liu *et al*. 2024), possibly reflecting variation among accessions and annotation methods. Importantly, AFL SRA 1002, from the Corsican citrus collection, is a confirmed interspecific hybrid, whereas Liu *et al*. (2024) studied a pure accession from the Givaudan Citrus Variety Collection at the University of California Riverside. The enrichment of tandem duplications within the CNL and NL classes suggests ongoing diversification of defense loci, making these clusters good candidates for future functional studies, particularly in relation to *Candidatus Liberibacter asiaticus* response.

While recent efforts have produced high-quality genome assemblies for Oceanian lime species (Nakandala, Furtado and Henry 2025), our study builds on these resources to address diversity and lineage structure at the population scale. Using the AFL genome as a reference, we resequenced other 54 Oceanian citrus accessions. These new data were joined with 78 previously published WGS resequencing data to perform a variant calling identifying close to 42 million SNPs. Among them a subset of more than 1.3 million high-confidence SNPs was used to establish a high-resolution phylogenomic reconstruction, resolving lineage structure across Oceanian citrus. While previous studies based on plastomes and limited nuclear markers (Nakandala *et al*. 2023a) supported the monophyly of Australian citrus, our use of whole-genome resequencing and availability of Papua New Guinea and New Caledonia species significantly expands this perspective. We demonstrate the monophyly of all Oceanian species and identified 84,678 DSNPs that fully differentiated all Oceanian species from all Asian ones.

*C. polyandra* appears as the basal species of the Oceanian clade. From chemotaxonomy (Berhow, Hasegawa and Manners 2000) proposed that *C. polyandra* may be a *C. japonica* Thunb. (1780) hybrid. The strong homozygosity of *C. polyandra* and the important differentiation of *C. polyandra* from all other *Citrus* species definitively discard this hypothesis.

The two species of the former *Oxanthera* genus cluster together constituting a clade with a large differentiation with other Oceanian species. As *C. polyandra*, they cluster basal to the remaining Oceanian lineages, confirming their inclusion in the *Citrus* genus as true species and reinforcing their unique evolutionary trajectory.

Interestingly, all the species from the former *Microcitrus* genus (Swingle and Reece, 1967) constitute a monophyletic group differentiated from the *C. glauca* clade (former *Eremocitrus* genus). This differentiation of *C. glauca* is consistent with an earlier evolutionary divergence and adaptation to a significantly harsher climate. *C. glauca* is also known for its strict resistance to HLB (Alves *et al*. 2021). Within the *Microcitrus* group we reveal the monophyletic origin of the three Papua-New Guinea species (*C. wintersii*, *C. warburgiana* and *C. wakonai*). The close relationship between *C. warburgiana* and *C. wakonai*, a new species endemic of Goodenough Island, recently discovered by Forster and Smith (2010) is observed both in nuclear and chloroplast phylogeny; however the high homozygosity of *C. wakonai* and the identification of DSNPs for *C. wakonai* and *C. warburgiana* all along the genome validate *C. wakonai* as an independent species. The Papua New Guinea clusters with a group of Australian species including *C. gracilis*, *C. australis* and *C. garrawayi* and the three unclassified “Cape York” accessions. We confirm the relatedness of *C.* sp Cape York accessions and *C. garrawayi* already observed by (Nakandala *et al*. 2023a) while the important homozygosity and the identification of DSNPs all along the genome support the “*C.* sp Cape York” group, that as a distinct morphology than *C. garrawayi*, as a new species. *C. australasica* clade which is of considerable commercial interest due to its described resistance/tolerance to HLB and culinary value and *C. inodora* clade constitute a second big cluster of the species of the former *Microcitrus* genus.

From resequencing data of the 103 representatives of ancestral species, we developed a robust panel of 4,390,992 diagnostic SNPs (DSNPs) capable of tracing the genomic origin of any citrus accession, including hybrids. This panel is effective to distinguish respectively 12 and 8 Oceanian species.

This tool allows precise clade assignment and admixture detection, with broad applications in germplasm curation, taxonomic clarification, and breeding program design as we have illustrated with the identification of admixed accessions in the germplasm collections of the partners and admixed hybrids from uncontrolled crosses. The identification of interspecific F1 hybrids and admixed accessions between Oceanian species in germplasm collections suggest a huge potential of interspecific hybridization between these monoembryonic non apomictic species. Many of these hybridizations may have occurred ex-situ during germplasm exchanges or propagation by seeds, but the *in-situ* vicinity of species may have resulted in wild interspecific hybridization. It may be the case for the introgression of *C. australis* in finger limes as, for example, wild accessions of both species have been described in close locations between Brisbane and Gold Coast.

Unlike traditional DNA markers, the DSNP panel enables unambiguous assignment of accessions to their ancestral clades all along the genome using genome-wide allelic signals. Beyond refining phylogenetic resolution, this approach enables the accurate identification of hybrid ancestries and varietal origins across global citrus, offering a transformative framework for germplasm classification and utilization. It is also a highly valuable tool for GWAS analysis based on ancestral haplotype and the driving of interspecific introgression schemes. This scalable resource represents a major advance for citrus diversity research and applied breeding.

Gene presence/absence analysis identified 1,090 genes absent in all Asian citrus accessions. These absences in Asian citrus likely result from gene loss during evolutionary divergence, or alternatively from the differentiation of ancestral gene content in Oceanian lineages. Among these, 127 genes were conserved across both Australian, New Caledonian and Papua New Guinean species, and 52 were unique to *C. australasica*. We acknowledge that some of the detected absences could also result from high sequence divergence or mapping bias during read alignment. Nevertheless, the identification of Oceanian-specific genes offers promising leads for understanding stress adaptation and immunity in wild citrus. These represent candidate loci potentially involved in stress tolerance, disease resistance, or adaptation to diverse ecological conditions.

Nevertheless, the AFL SRA 1002 genome is integrated into the Citrus Genome Hub (https://citrus-genome-hub.southgreen.fr/), which provides a genome browser and several user-friendly tools for molecular analyses.

Beyond phylogenetics, the SNP dataset generated provides a foundation for genome-wide association studies, selection scans, and trait mapping. Combined with the AFL reference genome, this framework represents the most comprehensive genomic toolkit to date for Oceanian citrus. It enables the strategic integration of wild diversity into breeding pipelines, particularly for traits like HLB resistance, environmental resilience, and fruit quality, and will support both fundamental research and global citrus improvement efforts.

## Conclusion

This study presents a high-quality, chromosome-scale genome assembly of *Citrus australasica* AFL SRA 1002 and the most extensive genomic survey to date of Oceanian citrus diversity. By integrating *de novo* assembly, genome annotation, and high-depth resequencing data of 132 accessions, we provide a robust framework for understanding the evolutionary history, structural variation, and gene content of this underexplored citrus lineage. The validation of novel species, the development of species diagnostic SNPs, and the identification of lineage-specific genes collectively underscore the value of Oceanian citrus as a reservoir of adaptive diversity. These resources lay the groundwork for functional, evolutionary studies, and targeted breeding strategies aimed at enhancing stress tolerance, disease resistance, and fruit quality in cultivated citrus. As global threats such as HLB intensify, the integration of wild diversity through genomics-informed approaches will be essential for sustaining citrus production and driving innovation in the years to come.

## Materials and Methods

### Genomic sequences of citrus accessions

#### Accession used for sequencing and optical mapping for *de novo* assembly

The genome assembly of the AFL was generated from accession SRA 1002, maintained in the Corsican field collection of the Citrus Biological Resource Center (Citrus BCR, INRAE-Cirad, San Giuliano, Corsica, France; Luro *et al*. 2017).

#### Resequencing of citrus accessions

For the diversity analyses, 54 *Citrus* accessions were newly resequenced and combined with 78 publicly available datasets from NCBI and EBI (Supplementary Table 2). Among the 54 new accessions respectively 18 were provided by the Bundaberg research station (Queensland Australia), 14 by Fundecitrus (Sao Paulo state, Brazil), 11 by Cirad-INRAE breeding program (Corsica, France), 6 by IVIA (Valencia, Spain), 3 by the Corsican field collection hosted by the Citrus BRC INRAE-Cirad (Corsica, France) and 2 by IAC (New Caledonia).

Genomic DNA was extracted using the DNeasy Plant Mini Kit (Qiagen). Two library types were used: (i) standard PCR-free Illumina libraries prepared for 33 accessions and sequenced on a HiSeq 4000 platform, and (ii) Tell-Seq linked-read libraries prepared for 21 accessions and sequenced on an Illumina NovaSeq platform, both with an average coverage of ∼30× per accession. R1 and R2 reads from Tell-Seq datasets were used for variant calling.

Demultiplexing, quality filtering, and adapter removal followed standard Genoscope and Montpellier GenomiX protocols. All datasets were processed using the same downstream pipeline to ensure comparability across sequencing technologies.

### Variant calling from resequencing data

Variant calling was performed using the Snakemake pipeline developed by Droc et al. (2024; https://gitlab.cirad.fr/agap/workflows/GATK), which automates GATK4 best-practice workflows for large-scale genomic analyses. Cleaned reads (fastp v0.20.1) from 54 newly sequenced accessions were combined with 78 publicly available datasets from EBI and NCBI (Supplementary Table 5). Reads were aligned to the new finger lime reference genome using BWA, and SNP/indel calling followed the GATK4 standard pipeline, including base quality recalibration and joint genotyping steps (GenomicsDBImport → GenotypeGVCFs). This unified workflow ensured consistent variant discovery and reproducibility across all datasets.

### Comparative genomic analysis and Gene family identification

To assess synteny and structural divergence between haplotypes, SyRI v1.3 (Goel *et al*. 2019) was used. Prior to this, the genome assemblies were aligned using nucmer from MUMmer v4.0.4 with the parameters --maxmatch -l 100 -c 500 -b 500, and filtered with delta-filter (-m -I 70 -l 100) (Marçais *et al*. 2018). For gene family identification, we used a comparative analysis to examine the conservation of gene repertoires among orthologs in the genomes of AFL SRA 1002, *C. australasica* (Singh *et al*. 2024) and *C. australis* (Nakandala *et al*. 2024a). First, we aligned all-to-all proteins using BLASTP (e-value of 1e−5). Genes were then clustered using OrthoMCL (1.4) implemented in OrthoVenn3 (Sun *et al*. 2023) with a Markov inflation index of 1.5 and a minimum e-value of 1e−15.

### Admixture analyses

Diagnostic SNPs (DSNPs) were identified following the two-step method of Ahmed et al. (2019) to discriminate ten Oceanian and seven Asian main clades, plus six species grouped within three composite clades. Analyses were based on 48 Oceanian and 55 Asian accessions (Supplementary Table 5). Briefly, SNPs with GST ≥ 0.5 were first used to detect introgressed regions, which were then excluded before a second run selecting fully diagnostic loci (GST = 1). Detailed procedures are provided in Ahmed *et al*. (2019) and Supplementary Note 2.

### Phylogenetic inference from nuclear and chloroplast SNPs

Maximum likelihood phylogenetic trees were reconstructed using IQ-TREE v2.2.6 based on aligned SNP datasets derived from nuclear and chloroplast genomes. The nuclear SNP alignment comprised 103 accessions and was built from a high-confidence set of concatenated loci, while the chloroplast SNP alignment included 100 accessions. For both datasets, the best-fit substitution model was selected using ModelFinder Plus (-m MFP), and branch support was assessed using 1000 ultrafast bootstrap replicates (-B 1000) and 1000 SH-aLRT replicates (-alrt 1000). The resulting trees were midpoint-rooted using Biopython v1.83 and exported in Newick format for downstream analyses and visualization.

### Gene loss analysis

Gene presence/absence variation (PAV) analysis was performed by aligning sequencing reads of each accession to the AFL reference genome with Bowtie2 v2.4.2 (“very-sensitive-local” mode).

PAVs were determined using SGSGeneLoss v0.1 (Golicz *et al*. 2015), classifying genes as present when ≥ 90 % of exons were covered by ≥ 5 reads (minCov = 5, lostCutoff = 0.9). Functional enrichment of gene sets (e.g., genes absent in Asian *Citrus*, conserved in Oceanian accessions, or specific to *C. australasica*) was performed in Cytoscape v3.10.3 using the BiNGO plugin (Maere *et al*. 2005) with FDR < 0.05, and visualized through EnrichmentMap (Merico *et al*. 2010) and AutoAnnotate (Kucera *et al*. 2016).

## Data Availability

Raw data from long-read sequencing and genome assemblies of *C. australasica* AFL SRA 1002 are available at EBI under the bioprojects PRJEB78475 and PRJEB78536. RNAseq data used for genome annotation is on EBI under the bioproject PRJEB104880. Raw resequencing data of the new 54 citrus accessions are available at EBI under the bioprojects PRJEB97093 (WGS) and PRJEB103773 (Tellseq).

## Acknowledgments

This work was supported by *France Génomique* (ANR-10-INBS-09–08; “Dynamo” project), European Feder-Guadeloupe region “Cavalbio” project, H2020 Innovation Action Program (grant #817526, preHLB project), French National Research Agency (grant ANR-23-CE20-0038, EpiHLB project), Ministry of Science and Innovation Spain (project PID2021-1281150R-I00), MCIN/AEI/10.13039/501100011033/FEDER, UE., and Hort. Innovation, Australia. The postdoctoral fellowship of LC is financed by the São Paulo Research Foundation (FAPESP), Brazil (#2024/09984-8). This work was supported by ISDM@Meso-Platform (University of Montpellier), CIRAD, and the bioinformatics group of UMR AGAP Institute, member of the IFB–South Green platform.

## Author Contributions

PO conceived and designed the study. BH and PO performed analyses and wrote the manuscript. MS, YF, MA, SL, and CM provided genetic resources. MM and PM carried out DNA/RNA extractions and RNA-Seq and TELL-Seq library preparations. CB, SD, CC, KL, BI, AL, PW, and JMA generated the genome sequencing and scaffolding of AFL SRA 1002. QP, AD, and GD produced the pseudochromosome assembly, genome annotation, and genome hub. MA led the genetic mapping and anchoring. KD and BH performed synteny analyses; BH conducted ortholog analyses. FSM and FM analyzed PRR genes; DG characterized repeated elements. PO and FC handled genotype calling; PO, AD, MA, LC, and FC identified DSNPs and performed admixture analyses. LC and MA analyzed gene loss. HB, FC, and PO prepared figures and tables and revised the manuscript. AL, PW, and JMA coordinated activities at Genoscope; NW at Fundecitrus; and PO, BH, and RM at CIRAD. PO, BH, RM, LP, NW, and AL contributed to funding acquisition.

## Conflict of Interest Statement

The authors declare no conflict of interest.

## Notes

### Competing Interest Statement

The authors have declared no competing interest.

