## Supplementary Material for "Chromosome-Scale Genome Assembly of Australian Finger Lime and Resequencing Reveal the Hidden Diversity of Oceanian Citrus"

### Supplementary Note 1: *De novo* AFL 1002 genome assembly and annotation

#### *De novo* nuclear genome assembly

The *de novo* genome assembly was obtained by combining nanopore long-read sequencing, Illumina PCR-Free sequencing and bionano optical mapping. The final pseudochromosome assembly was guided by Dotplot with *C. reticulata* CITRE genome assembly (Droc *et al.* 2024) and integration with an Oceanian citrus hybrid genetic map.

##### *PromethION library preparation and sequencing*

The libraries were prepared according to the '1D Genomic DNA by ligation (Kit 9 chemistry)' protocol, provided by Oxford Nanopore. Genomic DNA was first repaired and end-prepped with the NEBNext FFPE Repair Mix (New England Biolabs, Ipswich, MA, USA) and the NEBNext® Ultra™ II End Repair/dA-Tailing Module (NEB). DNA was then purified with AMPure XP beads (Beckmann Coulter, Brea, CA, USA), and sequencing adapters provided by Oxford Nanopore Technologies were ligated using Concentrated T4 DNA Ligase 2M U/ml (NEB). After purification with AMPure XP beads (Beckman Coulter) using dilution buffer (ONT) and wash buffer (ONT), the library was mixed with the sequencing buffer (ONT) and the Library Loading Bead (ONT) and loaded on the PromethION R9.4.1 Flow Cells. The nanopore long reads were not cleaned, and raw reads were used for genome assembly.

##### *Illumina PCR-Free library preparation and sequencing*

PCR free libraries were prepared using the Kapa Hyper Prep Kit (Roche, Basel, Switzerland). Briefly, genomic DNA (1-1.5µg) was sonicated using a Covaris E220 sonicator (Covaris, Woburn, MA, USA). Fragments were end-repaired and 3'-adenylated, and Illumina adapters (Bioo Scientific, Austin, TX, USA) were ligated following the manufacturer's instructions. Ligation products were purified twice with AMPure XP beads (Beckman Coulter Genomics,

Danvers, MA, USA) and quantified by qPCR (MxPro, Agilent Technologies, Santa Clara, CA, USA) using the KAPA Library Quantification Kit for Illumina Libraries (Roche). The library profile was assessed using an Agilent High Sensitivity DNA kit on the Agilent 2100 Bioanalyser. Libraries were paired-end sequenced on an Illumina HiSeq2500 instrument (Illumina, San Diego, CA, USA) using 250 base-length read chemistry. After the Illumina sequencing, an in-house quality control process was applied to the reads that passed the Illumina quality filters. The first step discards low-quality nucleotides (Q < 20) from both ends of the reads. Next, Illumina sequencing adapters and primer sequences were removed from the reads. Then, reads shorter than 30 nucleotides after trimming were discarded. These trimming and removal steps were completed using in-house-designed software based on the FastX package (Engelen and Aury; https://www.genoscope.cns.fr/fastxtend). The last step is to identify and discard read pairs that are mapped to the phage phiX genome, using SOAP aligner (Li *et al.* 2009) , and the Enterobacteria phage PhiX174 reference sequence (GenBank: NC_001422.1). This processing, described (Alberti *et al.* 2018), resulted in high-quality data.

***Optical mapping***

The direct label and stain labelling (using the DLE-1 enzyme) and the nick label repair and stain (NLRS) labelling (using the BspQI enzyme) protocols and Bionano Genomics protocols were followed with 750ng and 600ng of DNA, respectively. The chip loadings were performed as Bionano Genomics recommended.

***Long-read-based genome assembly***

First, we estimated genome size and heterozygosity rate using Genomescope2.0 (Ranallo-Benavidez, Jaron and Schatz 2020) with the Illumina reads as inputs. Using these values, we launched the Necat assembler (Chen *et al.* 2021) with the appropriate genome size and with all

Nanopore reads as inputs. Each assembly was then polished two times with HAPO-G (Aury and Istace 2021) and Illumina reads.

##### *Integration of the AFL genome assembly and Oceanian citrus genetic map*

Previously published GBS data of 171 'Fortune' mandarin × (*C. australis* × *C. inodora*) hybrids (Ollitrault *et al.* 2024) were used for variant calling using the new AFL genome assembly in pseudochromosomes as template. Detailed information for GBS analysis using ApeK I (Elshire *et al.* 2011) variant calling with the VCF-Hunter 2.1.0 pipeline (https://github.com/SouthGreenPlatform/VcfHunter; (Baurens *et al.* 2019), and genetic mapping with JoinMap5 (https://www.kyazma.nl/index.php/JoinMap/) can be found in Ollitrault *et al*. (2024).

##### *Scaffold sequence reconstruction*

Hybrid scaffolding: the BspQI and DLE-1 preparations were run on a single flow cell each. The generated molecules were assembled to produce optical maps using software provided by Bionano Genomics with the following two options: 'add pre-assembly' and non-haplotype without extend and split' (bionano solve and tools Version: 3.3_10252018). The two optical maps and the ONT contigs were subjected to the 2-enzyme hybrid scaffolding pipeline to generate the hybrid scaffolds. BisCoT (Istace, Belser and Aury 2020) was used to correct artifactual duplications (negative gaps) introduced during the scaffolding process.

##### *Dotplot analysis*

Comparisons between genome assemblies was performed using dotplot analysis with D-Genies (https://dgenies.toulouse.inra.fr/) with the Minimap2 v2.28 and "many repeats" options.

##### *Pseudochromosome construction*

The final genome assembly in pseudochromosomes was guided by dotplot analysis using the *C. reticulata* CITRE genome assembly in pseudochromosome as template (Droc *et al.* 2024) and the integration of the genetic map of the interspecific Oceanian hybrid *C. australis* × *C. australasica*. Chromosome numbering and orientation are identical to those of *C. clementina* V1.0 (https://phytozome-next.jgi.doe.gov/info/Cclementina_v1_0) and P. trifoliata V1.3.1 (https://phytozome-next.jgi.doe.gov/info/Ptrifoliata_v1_3_1).

#### Annotation of the Australian finger lime assembly

##### *RNA-Seq experiment and transcriptome assembly*

Three replicates of young leaves, mature leaves, flowers and fruit pulp from the Australian finger lime SRA 1002 were harvested on adult trees of the Corsican field collection hosted by the Citrus Biological Resource Center (Citrus BCR (INRAE-Cirad, San Giuliano, Corsica, France, Luro *et al.* 2017). RNA was extracted using the NucleoMag® kit (Macherey-Nagel GmbH & Co.KU, Düren, Germany) with the Kingfisher automated system (KingFisher Flex Purification System, ThermoFisher Scientific, Waltham, MA) according to the manufacturer's protocol. RNA integrity and quality were checked on TapeStation, screentape D5000. RNAseq libraries were prepared to obtain labelled and matched sequences using UDI indexes, following the protocol and recommendations from the Illumina Truseq kit. The libraries' integrity and quality were checked on TapeStation, screentape D5000, and assayed by qPCR. An equimolar pool of the libraries was then built up using the TruSeq Stranded mRNA Sample Preparation kit from Illumina. Pair-end sequencing (2 × 150 nt) was performed on an Illumina HiSeq 4000 platform at Genewiz.

Clean RNA-seq reads, which had been filtered by Trimmomatic (v0.39), were mapped onto the AFL genome assembly using MapSplice (Wang *et al.* 2010) with default parameters. Then the bam files were used with trinity (Grabherr *et al.* 2011) with the option for genome-guided transcriptome assembly. The obtained SAM alignment files were then converted into binary .BAM files using the samtools suite (Li *et al.* 2009). The quality of the reference transcriptome was then evaluated with Busco (Simão *et al.* 2015).

##### *Structural and functional annotation of the AFL SRA 1002 Genome assembly*

Automatic gene prediction was performed using the EuGene Eukaryotic Pipeline (EGNEP v.1.5 with EuGene v.4.2a) and Augustus (Stanke and Morgenstern 2005). EuGene is an integrative gene finder software that is able to combine several sources of information to predict genes (Sallet, Gouzy and Schiex 2019). A first run of Eugene (version 4.2a), was used to mask the genome via the repeat element detection tools integrated in the pipeline. Data from RepBase were provided to Eugene for integration into its genome-wide repeat detection and masking pipeline. The masked sequence was then used for ab-initio prediction via Augustus, with the *A. thaliana* plant model, for the prediction of genes and their structures. The results, in the form of a GFF3 file were incorporated as component in a second run of Eugene via the AnnotaStruct plug-in to generate the finale structural annotation. In addition, transcriptomic and protein evidences were retrieved and provided as input to Eugene. For transcriptomic data we included the AFL SRA 1002 transcriptome assembled by Trinity (of which we kept the longest representative for each transcript) and sequences from different species of the genus Citrus for which we defined very stringent parameters regarding the percentage of coverage (80%) and identity (90%). At the protein level, sequences extracted from the Uniprot database of the taxon Viridiplantae were used. The functional annotation of AFL SRA 1002 genome was performed by assigning their associated generic GO terms with the Blast2GO program using BLAST results from UniProt (UniProt 2021) with E-value 1e-5.

##### *Intragenomic duplication analysis*

Intragenomic collinearity analysis was performed using the SynMap pipeline (Haug-Baltzell *et al.* 2017) in the CoGe platform (https://genomevolution.org) to detect segmental duplications in the AFL SRA 1002 genome. Gene pairs were identified using DAGChainer with a maximum distance of 40 genes between matches (-D 40) and a minimum of five colinear gene pairs per block (-A 5). Syntenic blocks were subsequently merged using a maximum gap of 1,000 genes between blocks (-Dm 1000). The resulting duplication data were filtered to retain blocks longer than 7.6 kb (95th percentile) for visualization and downstream analysis. Coordinates and gene content of duplicated segments were extracted for functional assessment.

***Annotation of PRR***

Protein sequences and corresponding GFF3 annotation files from the AFL genome were used as input data for downstream analyses. Domain annotation of all predicted proteins was performed using InterProScan v5.48-83.0 (Jones *et al.* 2014) with default parameters. The search integrated multiple domain databases, including COILS, Gene3D, HAMAP, MOBIDB, PANTHER, Pfam, PIRSF, PRINTS, ProDom, PROSITE, SFLD, SMART, SUPERFAMILY, and TIGRFAM. Results were exported in tab-separated values (TSV) format.

The identification and classification of pattern recognition receptors (PRRs) and resistance (R) proteins were carried out using RRGPredictor (Silva and Micheli 2020), which assigns each protein to one of 13 structural classes: T, TN, TNL, RPW8NL, RLP, RLK, RLKGNK2, N, NL, CN, CNL, MLO, and UNKNOWN. The UNKNOWN class was used solely for estimating the total number of PRR genes and was excluded from class-specific analyses. Chromosomal localization of genes was visualized using TBtools.

Gene duplication analysis was conducted with MCScanX (Wang *et al.* 2012) under default parameters. Duplication events were analyzed and visualized for each identified PRR and R gene class, generating both tabular outputs and graphical representations of gene collinearity and tandem duplication.

***Annotation of repeated sequences in the genome assembly***

The complete diversity of repeated sequences was characterized including the annotation of transposable elements (TEs), satellite DNAs (or tandem repeats), endogenous caulimovirids elements (ECVs) and Simple Sequence Repeats (SSRs). Repeated sequences were identified using the faster method developed by Giraud *et al.* (2025). First, the repeat-finding programs REPET (Quesneville *et al.* 2005; Flutre *et al.* 2011), MITE-Hunter (Han and Wessler 2010) and TAREAN (Novák *et al.* 2017) were used to construct a unique library of consensus sequences consisting of all TEs, ECVs and satellite DNAs identified de novo in the genome assembly. The current library was then compared with the citrus reference repeated sequence library, which was previously developed following the annotation of repeated sequences in 12 *Citrus* species (Giraud 2025). Using BLASTn, all consensus from the current library with a percentage of identity lower than 80% with the consensus of the citrus reference library were retained because they represented new diversity not already found in the reference library (<80% of identity on <80% of the smallest sequences, e-value <1e-06 (Camacho *et al.* 2009). This set of consensus was classified using PASTEC (Hoede *et al.* 2014) and sequences features were checked according to the method developed by (Giraud *et al.* 2025). Finally, the quantity of repeated sequences on the genome assembly and their localization were determined by aligning the citrus reference library enriched with the new consensus set on the pseudoChromosomes using TEannot, a pipeline included in REPET package. The density of the different repeated sequences along pseudoChr omosomes was calculated with a 1-Mb sliding window and represented on circular plot using circos v0.69.9 (Krzywinski *et al.* 2009) (Krzywinski *et al.* 2009).

#### Supplementary Note 2 : Identification of admixture along the four ancestral genomes

TraceAncestor software (Ahmed *et al.* 2019) was adapted for WGS data from diploid genotypes. Genotyping data were used instead of read counts to conduct maximum-likelihood tests of each ancestor contribution (i.e., allelic doses = 0, 0.5, 1 dose) in genomic windows of 20 DSNPs.

Considering A as the ancestral taxon allele at each locus, B as the alternative allele (allele from another ancestor), and e as the sequencing error rate (fixed at 0.01 for our analysis), the probability of observing i ancestral alleles and j alternative allele for the 20 DSNPs loci according to the tested model of ancestral doses for diploid plants was as follows:

$$2 \text{ ancestral doses (AA): } C \times P(A)^{i} \times P(B)^{j} = C \times (1\text{-}e)^{i} \times e^{j}$$

$$1 \text{ ancestral dose (AB): } C \times P(A)^{i} \times P(B)^{j} = C \times 0.5^{i} \times 0.5^{j}$$

$$0 \text{ ancestral dose (BB): } C \times P(A)^{i} \times P(B)^{j} = C \times e^{i} \times (1\text{-}e)^{j}$$

where C is a combinatorial coefficient which depends on the observed data but remains constant between the models; P(A) and P(B) respectively the theoretical frequencies of the ancestor and alternative alleles.

We compared one dose hypothesis (heterozygosity) with 0 and two dose hypotheses by calculating a LOD (logarithm of odds) score :

$$\text{LOD } (0/1) = \text{Log10 } ((e^{i} \times (1\text{-}e)^{j})/(\, 0.5^{i} \times 0.5^{j}))$$

$$\text{LOD } (1/2) = \text{Log10 } ((0.5^{i} \times 0.5^{j})/((1\text{-}e)^{i} \times e^{j}))$$

t being the selected LOD threshold value, the process to conclude for a dose was as follows:

If LOD (0/1) >= t : 0 ancestral taxon dose

If – t < LOD < t : undetermined dose

If LOD (0/1) <= - t and LOD (1/2) >= t : 1 ancestral taxon dose

If LOD (0/1) <= - t and –t < LOD (1/2) < t : undetermined dose

If LOD (0/1) <= - t and LOD (1/2) <= - t : 2 ancestral taxon doses

The t value (LOD threshold) was fixed at 3 for all analysis

During this step, the number and position of windows varied between the ancestral species according to the density and positions of the DSNPs of each of the considered species. To integrate the information obtained for all ancestral species, we sub-divided the genome into successive fragments of 100 kb. For each ancestor and for each 100 kb genomic fragment, the corresponding window of 20 DSNPs was identified and the ancestral dose of this window was attributed to the considered 100 kb genomic fragment. In cases in which the sum of ancestral contributions exceeded 2, the corresponding windows were considered undetermined for the two haplotypes. The identification of DSNPs and mosaic structure analyses were performed using the SniPlay web tool (https://sniplay.southgreen.fr/cgi-bin/home.cgi).

**Supplementary Table 1:** Summary statistics of the Australian finger lime (SRA 1002) genome scaffold assembly.

| Statistic | Value (Mbp or %) |
|---|---|
| Total number of scaffolds | 403 |
| Cumulative scaffold size | 450.8 Mbp |
| N50 scaffold size | 28.40 Mbp |
| Number of scaffolds ≥ N50 | 7 |
| N80 scaffold size | 5.41 Mbp |
| Number of scaffolds ≥ N80 | 19 |
| N90 scaffold size | 0.72 Mbp |
| Number of scaffolds ≥ N90 | 47 |
| Minimum scaffold size | 0.011 Mbp |
| Maximum scaffold size | 38.55 Mbp |
| Average scaffold size | 1.12 Mbp |
| auN (area under the Nx curve) | 21.14 Mbp |
| % GC content | 34.72% |
| % N bases | 6.48% |

**Supplementary Table 2.** *C. australis* × *C. inodora* genetic map: distribution of the SNP markers among linkage groups and AFL pseudochromosomes

| LG\SC | 1 | 2 | 3 | 4 | 5 | 6 | 7 | 8 | 9 | Mks/LG | LG size cM |
|---|---|---|---|---|---|---|---|---|---|---|---|
| 1 | 266 | 0 | 0 | 0 | 0 | 0 | 0 | 0 | 0 | 266 | 106.28 |
| 2 | 0 | 313 | 0 | 0 | 0 | 0 | 0 | 0 | 0 | 313 | 123.47 |
| 3 | 0 | 0 | 469 | 0 | 0 | 0 | 0 | 0 | 0 | 469 | 155.44 |
| 4 | 0 | 0 | 0 | 338 | 0 | 0 | 0 | 0 | 0 | 338 | 103.82 |
| 5 | 0 | 1 | 0 | 0 | 302 | 0 | 1 | 0 | 2 | 306 | 94.03 |
| 6 | 0 | 0 | 1 | 0 | 0 | 241 | 0 | 0 | 0 | 242 | 77.36 |
| 7 | 1 | 0 | 0 | 2 | 0 | 0 | 256 | 0 | 0 | 259 | 92.1 |
| 8 | 0 | 0 | 0 | 0 | 0 | 0 | 0 | 230 | 0 | 230 | 95.81 |
| 9 | 0 | 0 | 1 | 0 | 0 | 0 | 0 | 0 | 198 | 199 | 95.3 |
| Mks /Sc | 267 | 314 | 471 | 340 | 302 | 241 | 257 | 230 | 200 | 2,622 | 943.61 |

**Supplementary Table 3 :** Results of the *AFL SRA 1002* reference transcriptome implementation

| Sample | Sequencing Volume (Gb) | Nb of transcripts | BUSCO score |
|---|---|---|---|
| Flowers | 67.8 | 76 534 | 95 |
| Fruit pulp | 23 | 43 806 | 76 |
| Mature leaves | 51.6 | 58 892 | 91 |
| Young leaves | 54 | 54 718 | 94 |

**Supplementary Table 4:** Classification of new repeated sequences identified on the AFL SRA 1002 genome assembly

| | | Number of consensus sequences | Number of copies (full / truncated) | Genome coverage (%) |
|---|---|---|---|---|
| **Retrotransposons (Class I elements)** | | | | |
| LTR *Copia* (RLC) | *Tork* | 4 | 4 / 191 | 0.051% |
| | *Retrofit* | 3 | 17 /17 | 0.037% |
| LTR *Gypsy* (RLG) | *Tat* | 1 | 0 / 6 | 0.000% |
| | *Crm* | 1 | 0 / 9 | 0.001% |
| LINEs (RIX) | | 6 | 9 / 902 | 0.067% |
| **DNA transposons (Class II elements)** | | | | |
| TIRs | *Mutator* (DTM) | 2 | 4 / 36 | 0.011% |
| | *Pif-Harbinger* (DTH) | 1 | 0 / 27 | 0.002% |
| | *Tc1-Mariner* (DTT) | 1 | 2 / 25 | 0.003% |
| Unclassified MITEs (DXX-MITE) | | 5 | 8 /11 | 0.000% |
| Helitron (DHX) | | 1 | 4 / 4 | 0.001% |
| **TOTAL Transposable Elements** | | **25** | **48 / 1128** | **0.173%** |

**Supplementary Table 5.** Number of duplication events of PRR receptors and R genes present in each class in AFl SRA 1002 genome assembly. Genes involved in two duplication events are highlighted in bold.

| Class | Quantity of duplication events | Duplication events |
|---|---|---|
| CN | 1 | Ciaus5g05200:Ciaus5g05670 |
| CNL | 4 | Ciaus5g14600:**Ciaus5g18130**<br>Ciaus5g17860:**Ciaus5g18130**<br>Ciaus5g16120:**Ciaus5g16660**<br>Ciaus5g16040:**Ciaus5g16660** |
| MLO | 1 | Ciaus5g40430:Ciaus7g00190 |
| N | 0 | - |
| NL | 4 | Ciaus5g01860:Ciaus5g02490<br>Ciaus5g02180:Ciaus5g02790<br>Ciaus5g14630:Ciaus5g17990<br>Ciaus5g05320:Ciaus5g05760 |
| RLK | 1 | Ciaus6g16950:Ciaus6g17410 |
| RLKGNK2 | 2 | Ciaus2g08190:**Ciaus4g25280**<br>Ciaus3g05430:**Ciaus4g25280** |
| RLP | 0 | - |
| RPW8NL | 0 | - |
| T | 2 | Ciaus2g02190:Ciaus2g02750<br>Ciaus5g08260:Ciaus5g08640 |
| TN | 0 | - |
| TNL | 3 | Ciaus3g34420:Ciaus3g35380<br>Ciaus3g30790:Ciaus3g31190<br>Ciaus5g08170:Ciaus5g08580 |

**Supplementary Table 7**: DSNPs of the species of the *C. wakonai* + *C. warburgiana C. garrawayi* + Unknown and *C. medica* + *C. indica* main clades

| | **Chr1** | **Chr2** | **Chr3** | **Chr4** | **Chr5** | **Chr6** | **Chr7** | **Chr8** | **Chr9** | **Total** |
|---|---|---|---|---|---|---|---|---|---|---|
| *C. wakonai* | 10535 | 11471 | 13261 | 9074 | 8202 | 8396 | 9157 | 7992 | 6938 | **85026** |
| *C. warburgiana* | 11079 | 16095 | 14531 | 7831 | 6405 | 9875 | 10307 | 8429 | 9255 | **93807** |
| Unknown | 12081 | 12407 | 16665 | 8244 | 9684 | 13664 | 8468 | 11747 | 7095 | **100055** |
| *C. garrawayi* | 7082 | 8952 | 10753 | 5168 | 6803 | 8731 | 5364 | 8184 | 4383 | **65420** |
| *C. medica* | 27681 | 28322 | 38403 | 27610 | 23379 | 25537 | 21649 | 24197 | 22345 | **239123** |
| *C. indica* | 15287 | 16623 | 22038 | 15205 | 12822 | 13589 | 12840 | 13619 | 11760 | **133783** |
| Total | 83745 | 93870 | 115651 | 73132 | 67295 | 79792 | 67785 | 74168 | 61776 | **717214** |

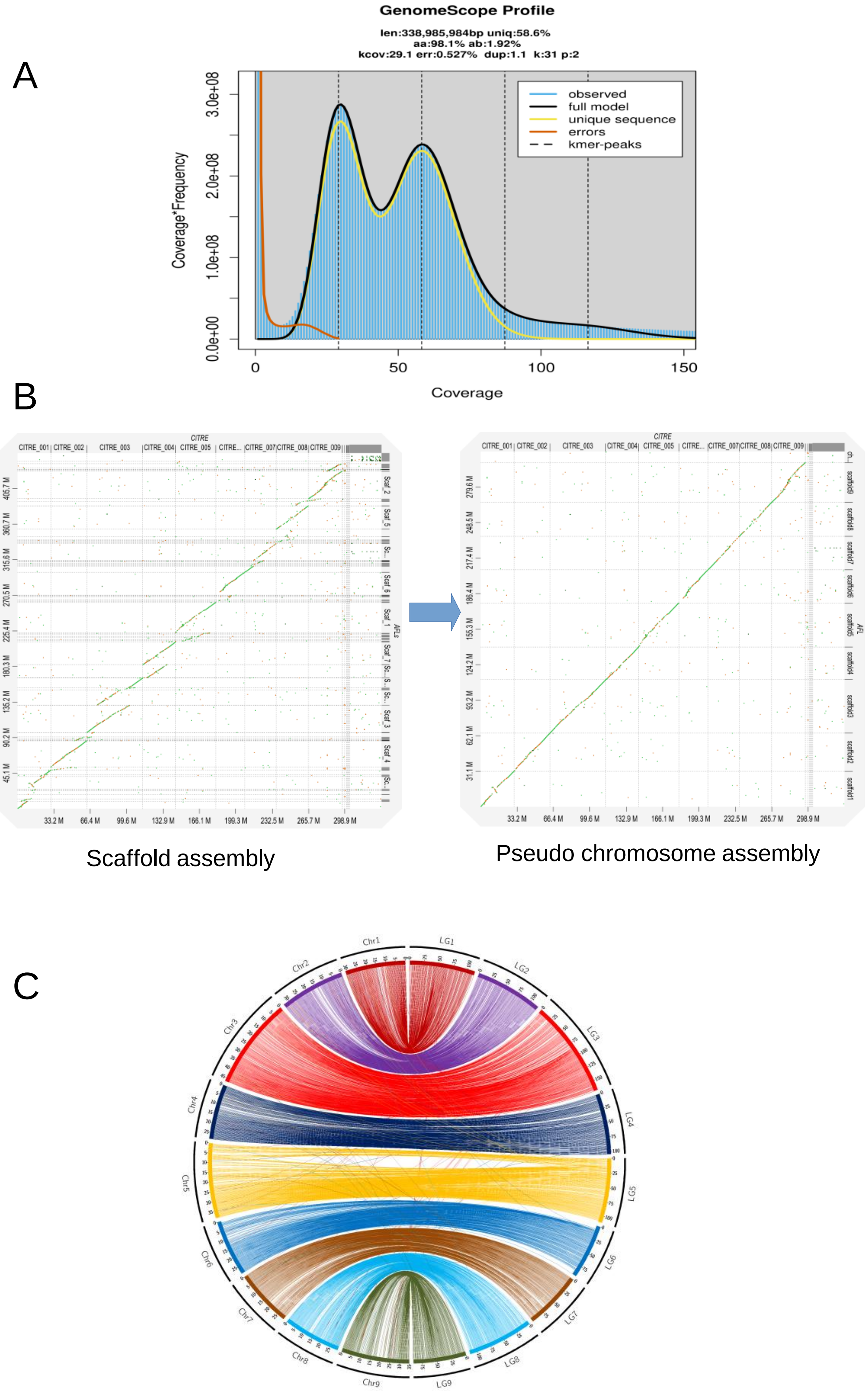


**Supplementary Figure 1: Interspecific mosaic structure of the sequenced accession and implication in duplicated areas on the genome assembly**. **A)** GenomeScope2.0 k-mer analysis based on Illumina short reads estimates a genome size of 338 Mb and a heterozygosity rate of 1.92%, supporting the hybrid nature of this accession. **B)** DotPlot of preliminary scaffold assembly (left) and final pseudochromosome scale assembly with the CITRE genome (Cleopatra mandarin; Droc *et al.* 2024). **C)** Anchorage of 7819 genes the *Citrus* consensus genetic map (Ollitrault *et al.* 2024) on the pseudochromosome assembly.

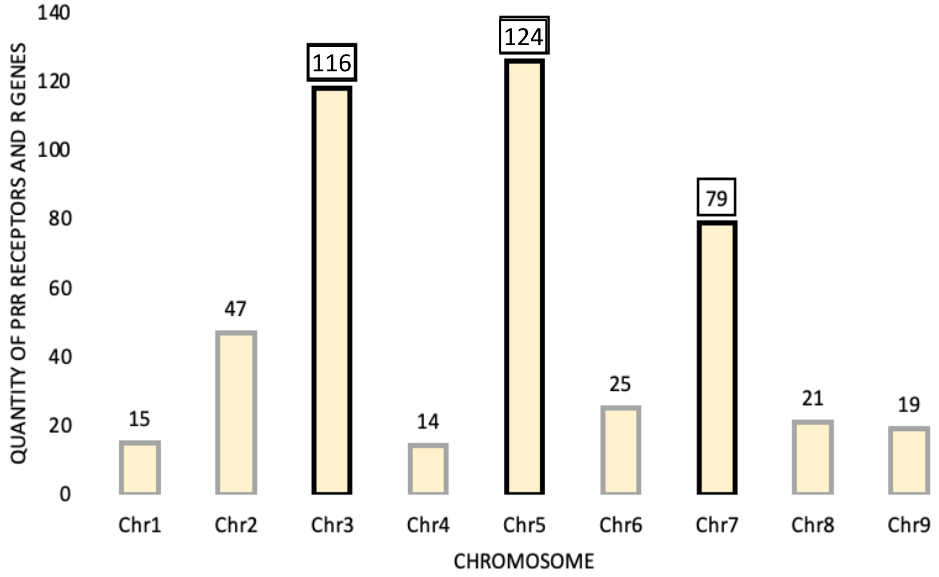


**Supplementary Figure 2:** Distribution of PRR receptors and R genes from the well-defined classes across the chromosomes of AFL SRA 1002.

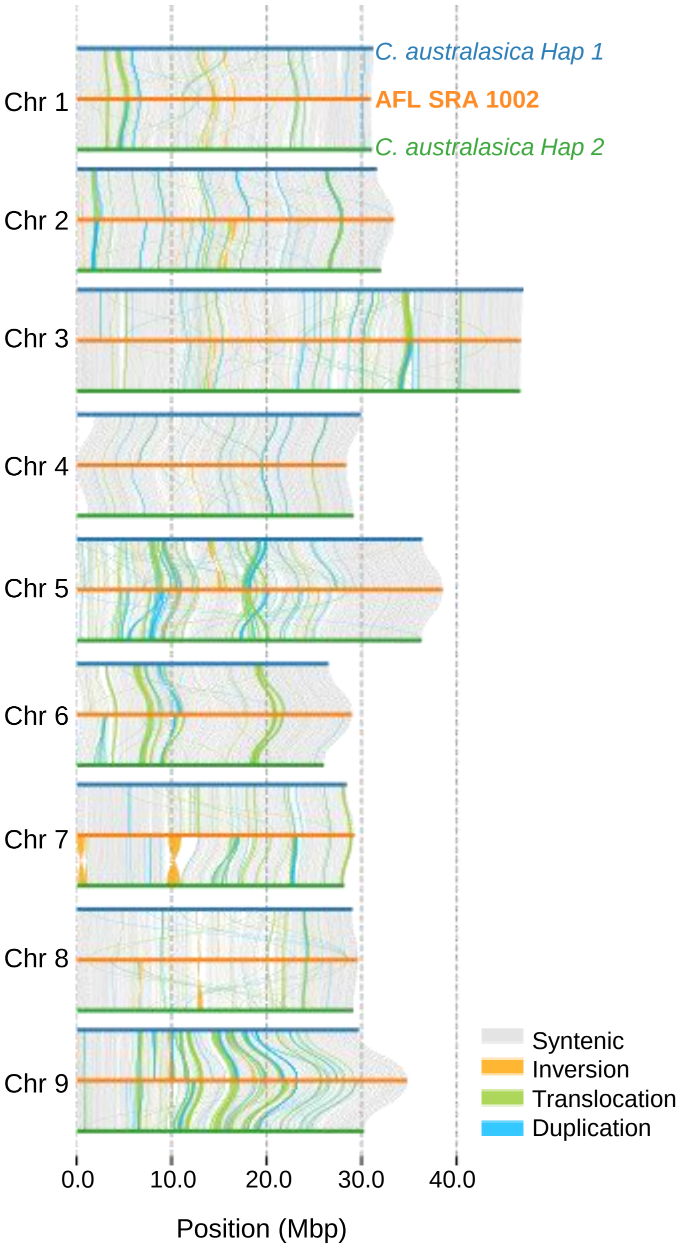


**Supplementary Figure 3:** Comparative structural genomics between AFL SRR 1002 and the two haplotypes of *Citrus australasica* assemblies as reported by Nakandala *et al*. (2024). Conserved syntenic blocks are depicted in grey, while regions lacking alignment are shown in white. Distinct categories of genomic rearrangements are color-coded according to their type. Structural variation was assessed using the Synteny and Rearrangement Identifier (SyRI) tool.

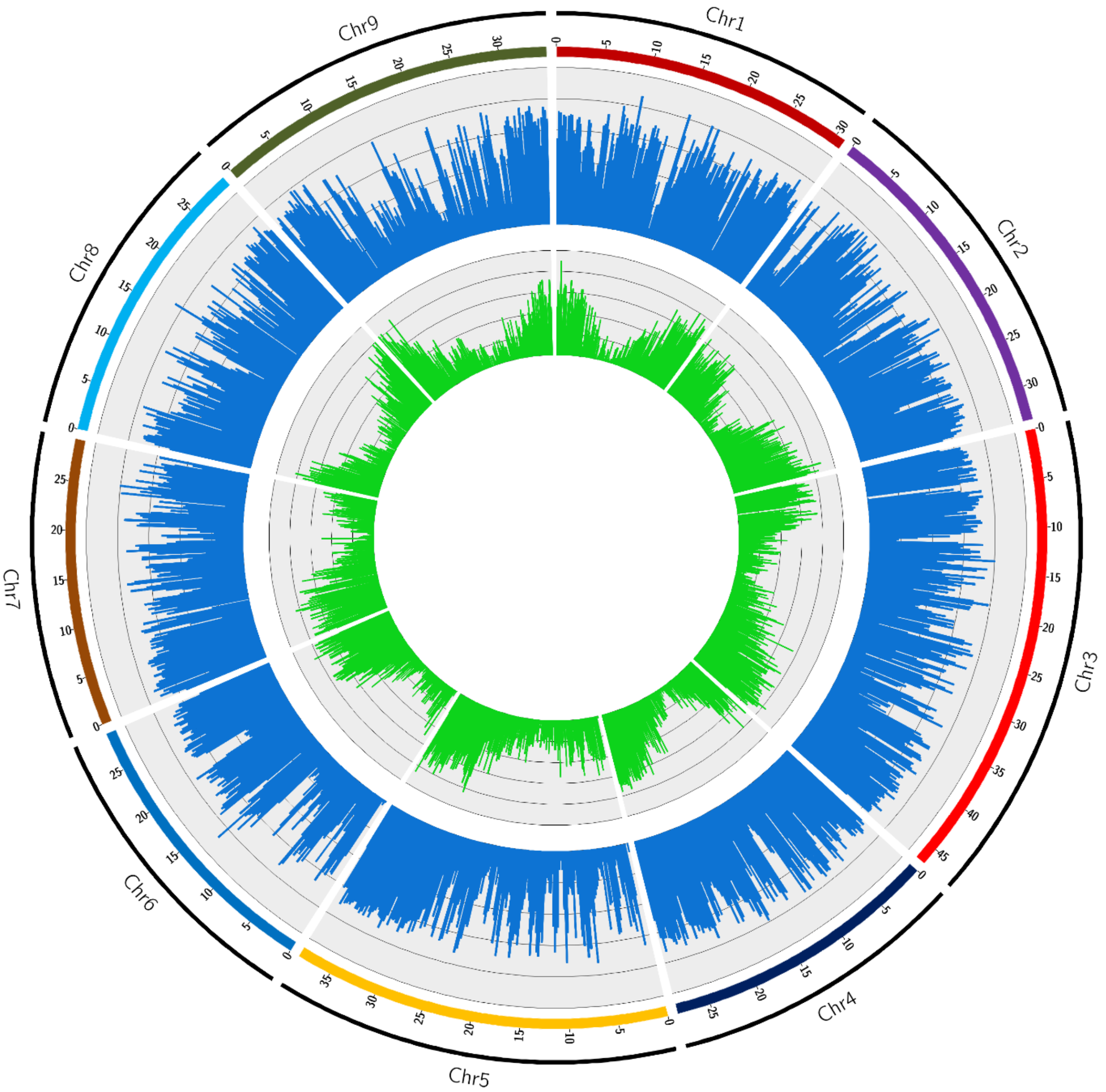


**Supplementary Figure 4:** Distribution of SNPs and genes along the AFL genome. External ring: SNP density; Windows of 200 kb; scale 0 to 50000 SNPs / 200 kb. Internal ring: Gene density Windows of 200 kb; scale 0 to 50 genes / 200 kb.

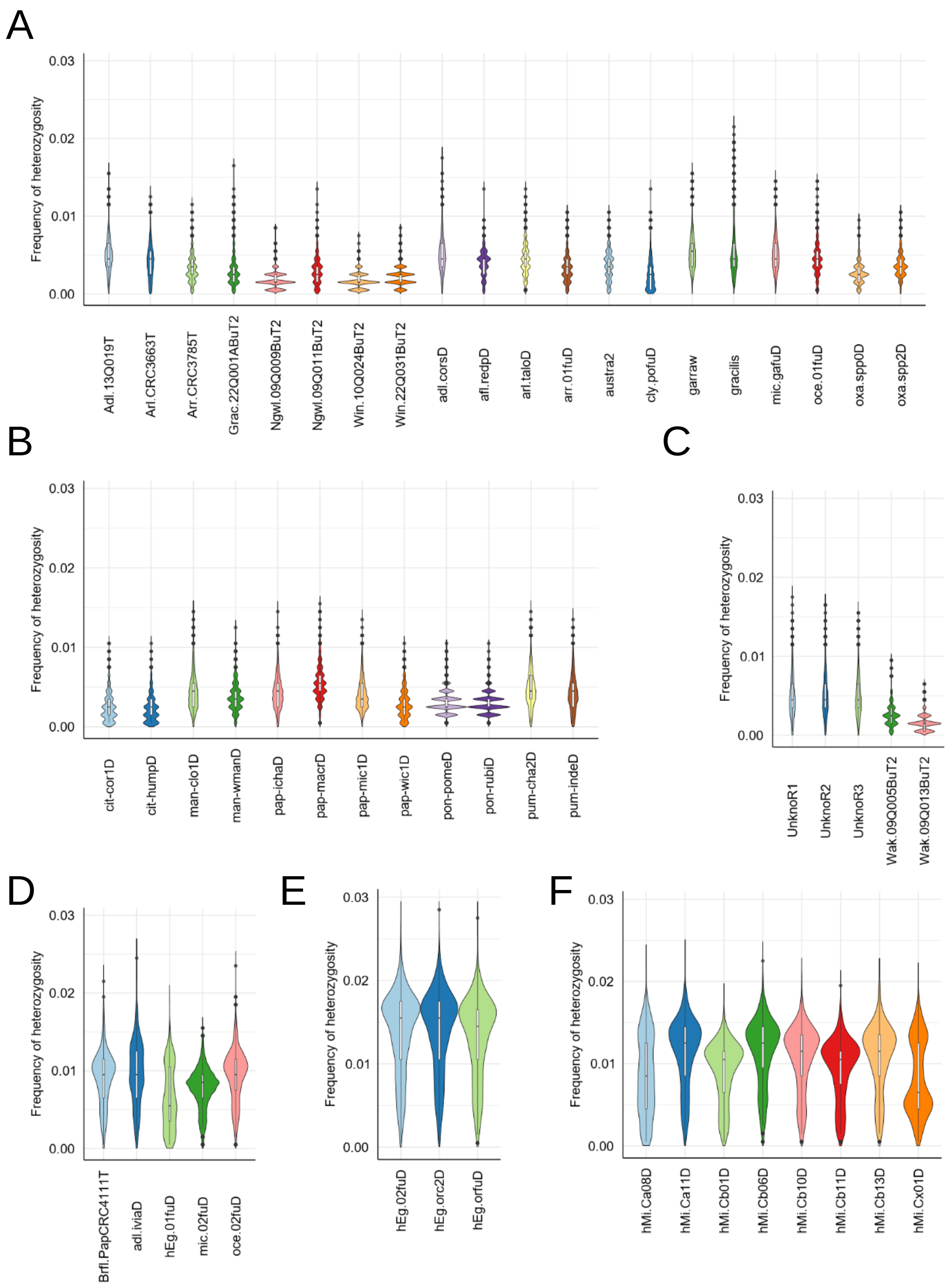


**Supplementary Figure 5: Heterozygosity value distribution (frequency of heterozygosity in successive 200 kb windows)**. **A)** Oceanian species representatives; **B)** Asian species representatives; **C)** Unknown Cape York sp. and *C. wakonai* representatives; **D)** Other interspecific Oceanian accessions; **E)** Eremorange accessions; F) Randomly admixed hybrids of the Cirad breeding project.

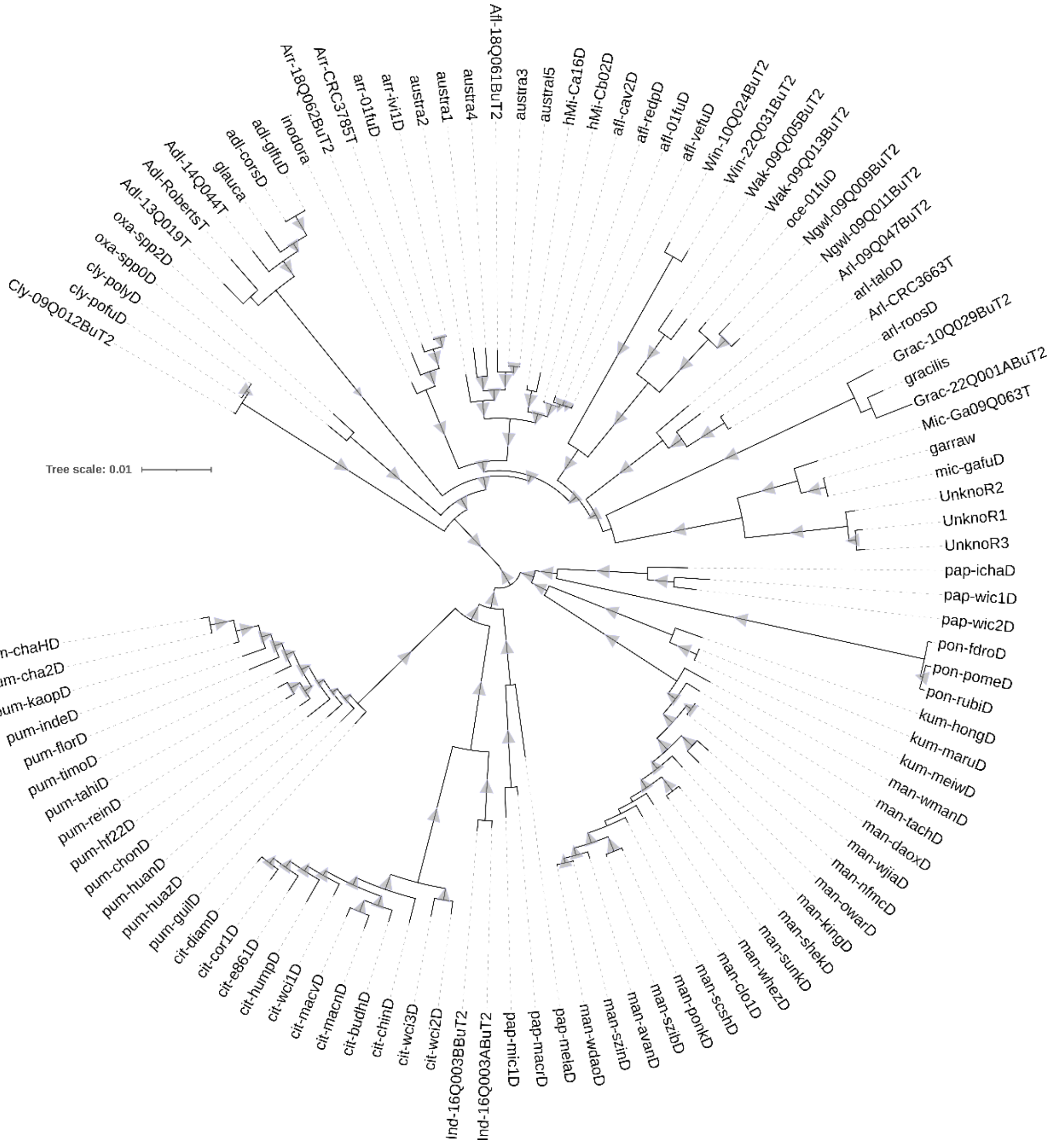


**Supplementary Figure 6:** Maximum likelihood phylogenetic tree based on 1,322,933 nuclear SNPs, with explicated codes for each genotype. Triangles indicate branch support (Boostrap >90; 1000 replicates), with size proportional to support values. The scale bar represents 0.01 substitutions per site. Genotypes list is available in Supplementary Table 5.

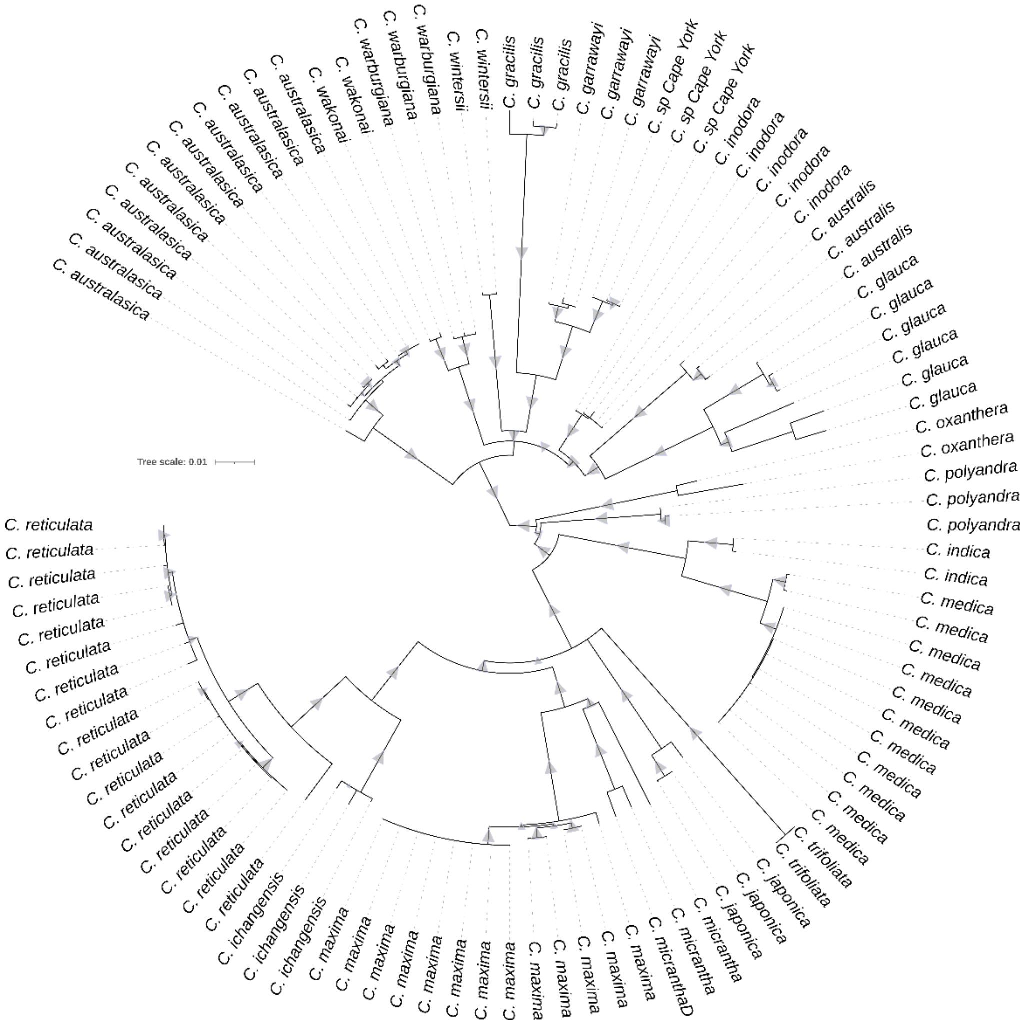


**Supplementary Figure 7:** Maximum likelihood phylogenetic tree based on 3,695 SNPs derived from the *C. sinensis* plastome, Triangles indicate branch support (Boostrap >90; 1000 replicates), with size proportional to support values. The scale bar represents 0.01 substitutions per site.

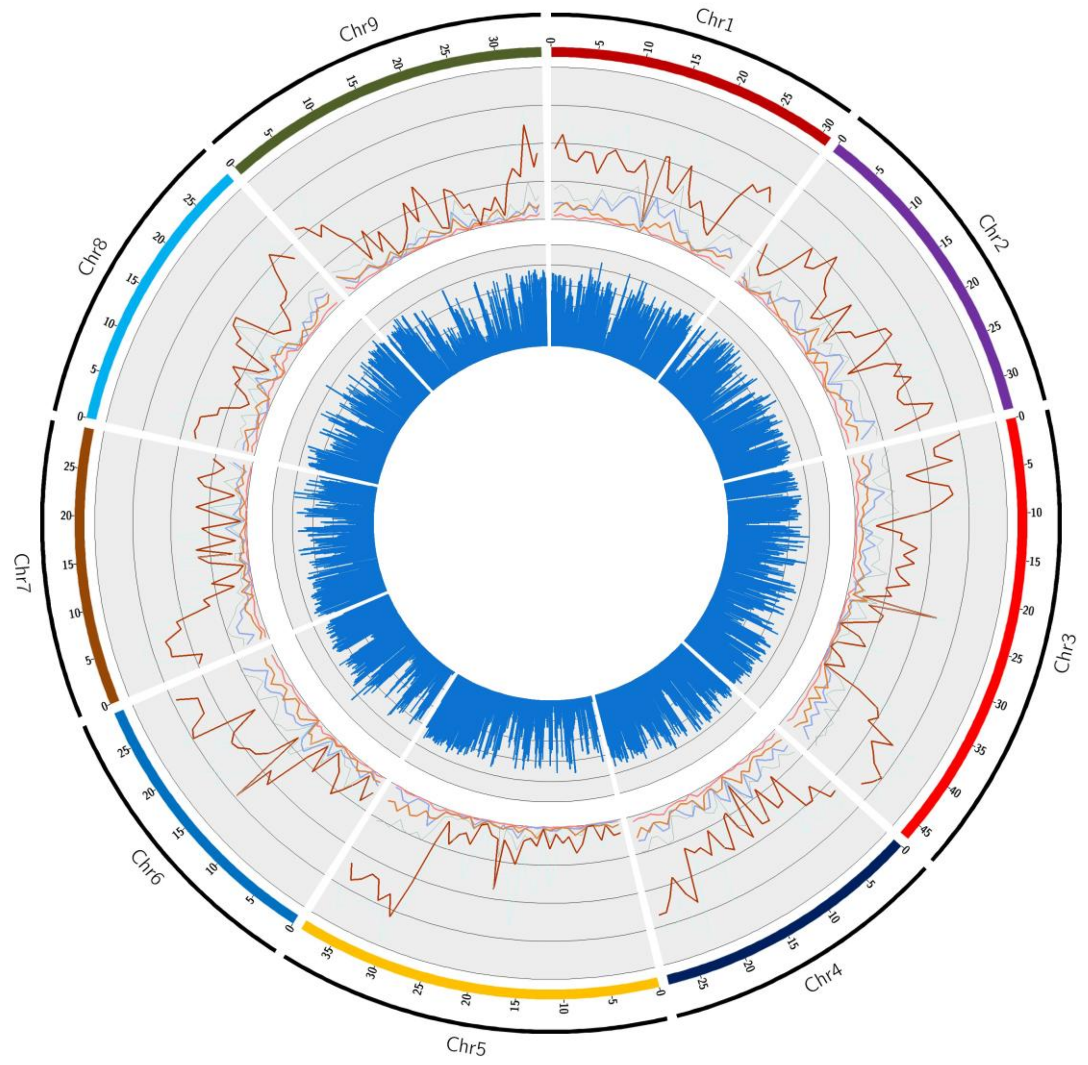


**Supplementary Figure 8:** Distribution along the genome of the DSNPs of Australian species. Internal ring: SNP density. Windows of 200 kb; scale 0 to 50000 SNPs / 200 kb. External ring: DSNP density; dark brown: *C. glauca*; light brown: *C. inodora*; salmon: *C. australasica*, light blue: *C. australis*; peacock blue: *C. garrawayi*; + Unknown Cape York sp.; sky-blue: *C. gracilis*. Windows of 1Mb; scale: 0 to 4000 DSNPs / Mb.

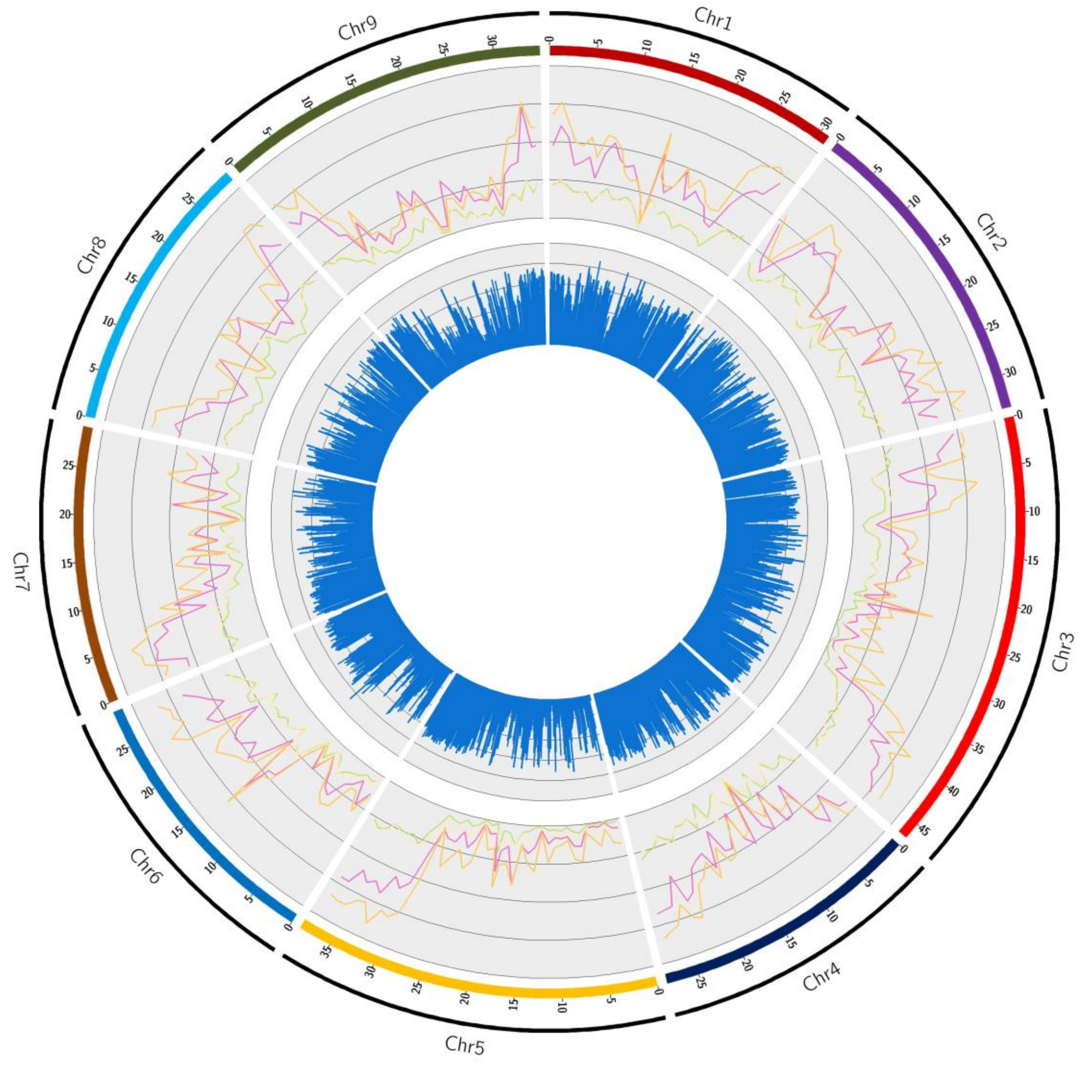


**Supplementary Figure 9:** Distribution along the genome of the DSNPs of Papua-New-Guinean and New-Caledonian species. Internal ring: SNP density. Windows of 200 kb; scale 0 to 50000 SNPs / 200 kb. External ring: DSNP density; pink: *C. wintersii*; yellow-green: *C. warburgiana* + *C. wakonai*; sunset yellow: *C. polyandra*; blond: *Oxanthera* sp. Windows of 1Mb; scale: 0 to 4000 DSNPs / Mb.

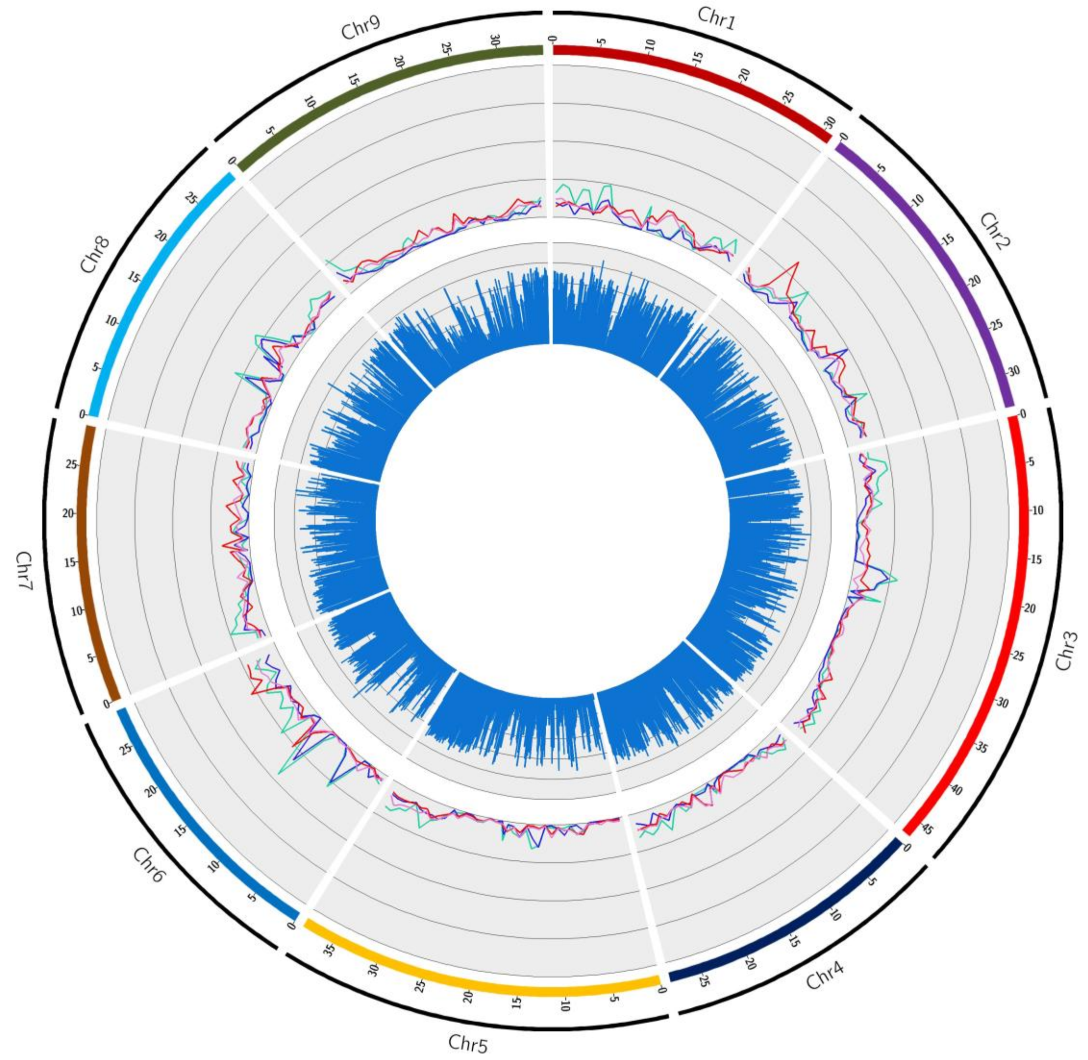


**Supplementary Figure 10**: Distribution along the genome of Oceanian DSNPs of the Oceanian subclades. Internal ring: SNP density. Windows of 200 kb; scale 0 to 50000 SNPs / 200 kb. External ring: DSNP density; green: Unknown Cape York sp.; blue: *C. garrawayi*; red: *C. warburgiana*; pink: *C. wakonai*. Windows of 1Mb; scale: 0 to 4000 DSNPs / Mb.

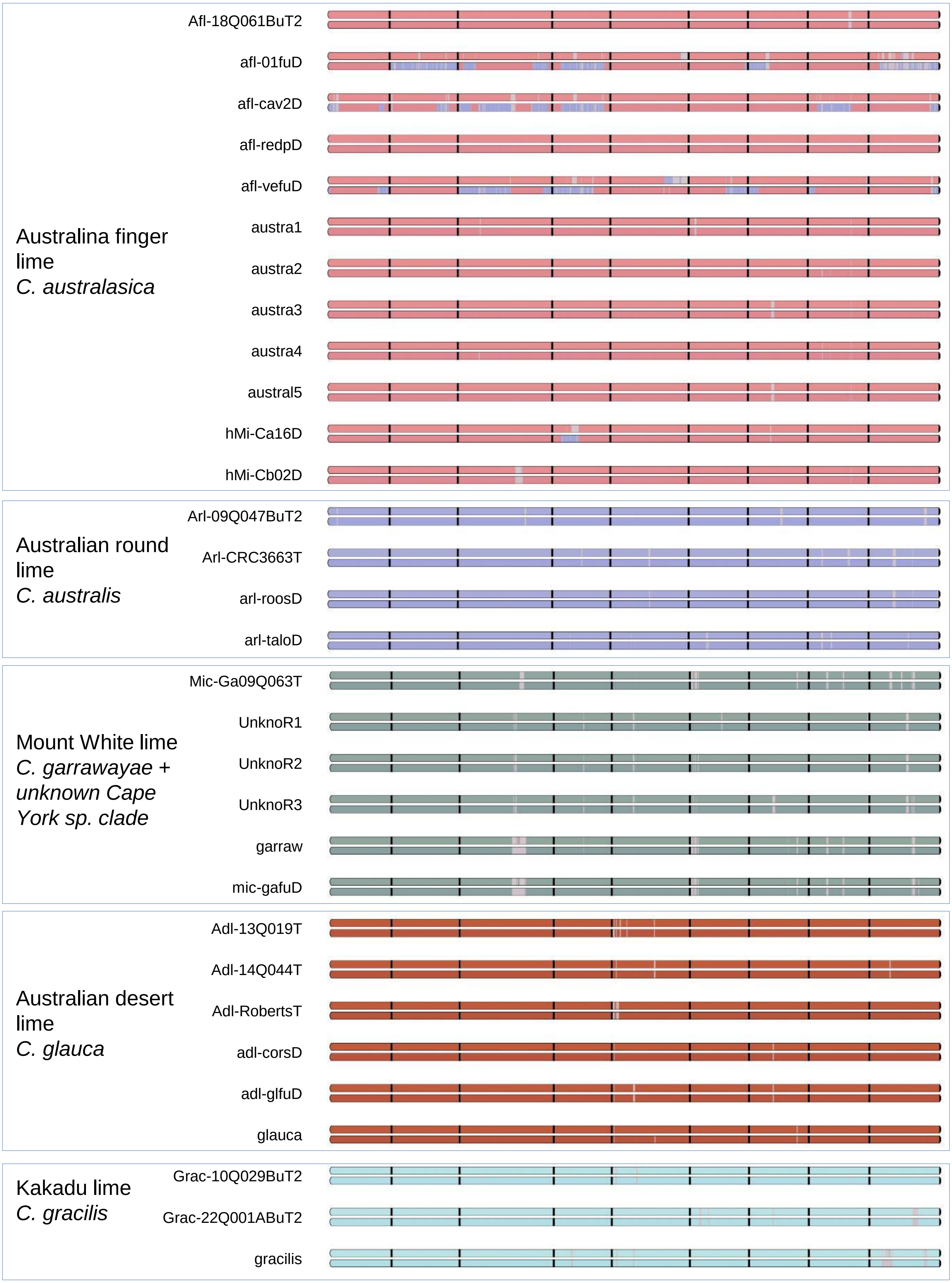


**Supplementary figure 11:** Phylogenomic mosaic structure along the nine chromosome (separated by black dash) of the accessions representatives of the main Oceanian clades (first part).

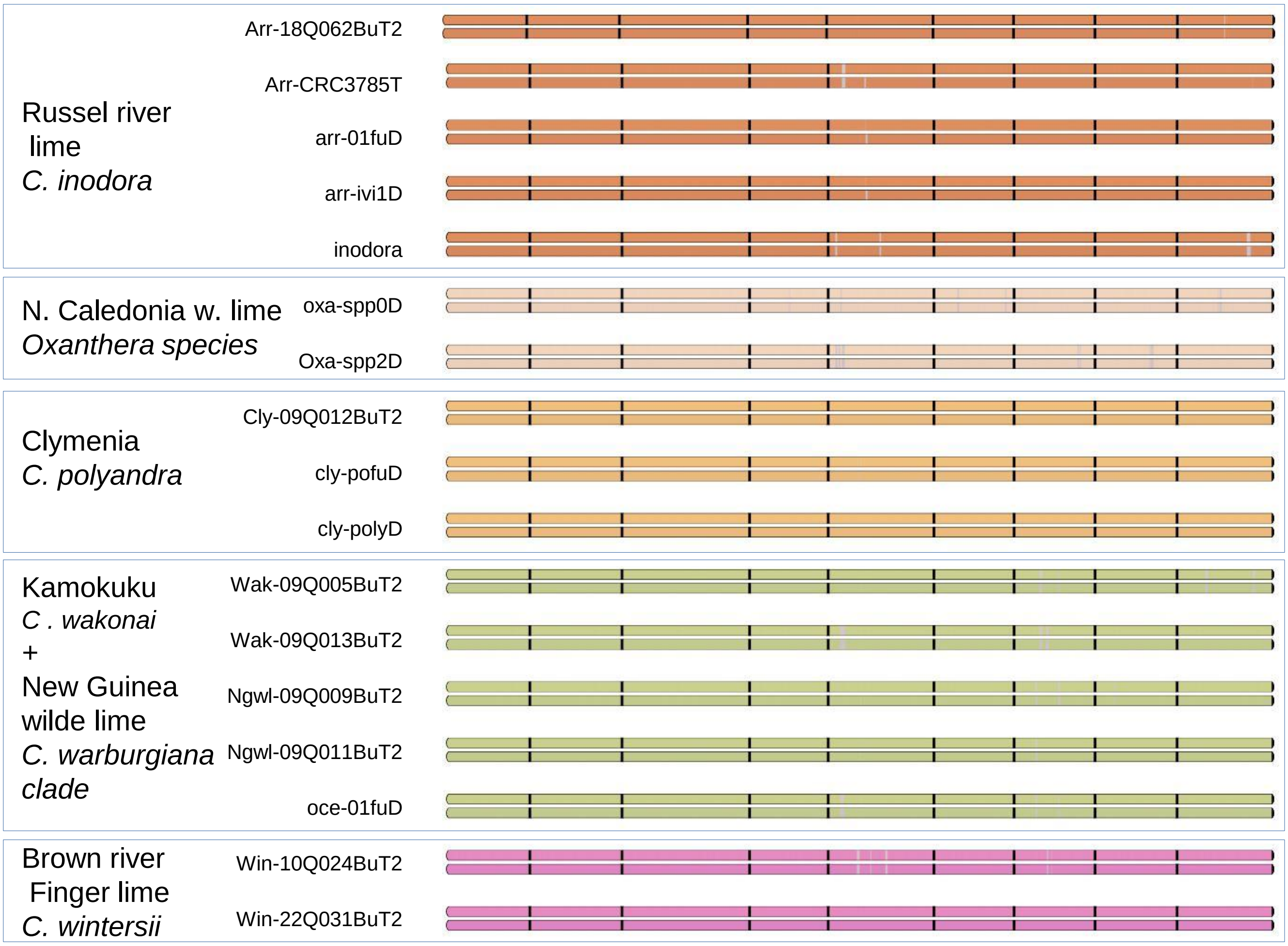


**Supplementary figure 11:** Phylogenomic mosaic structure along the nine chromosome (separated by black dash) of the accessions representatives of the main Oceanian clades (second part).

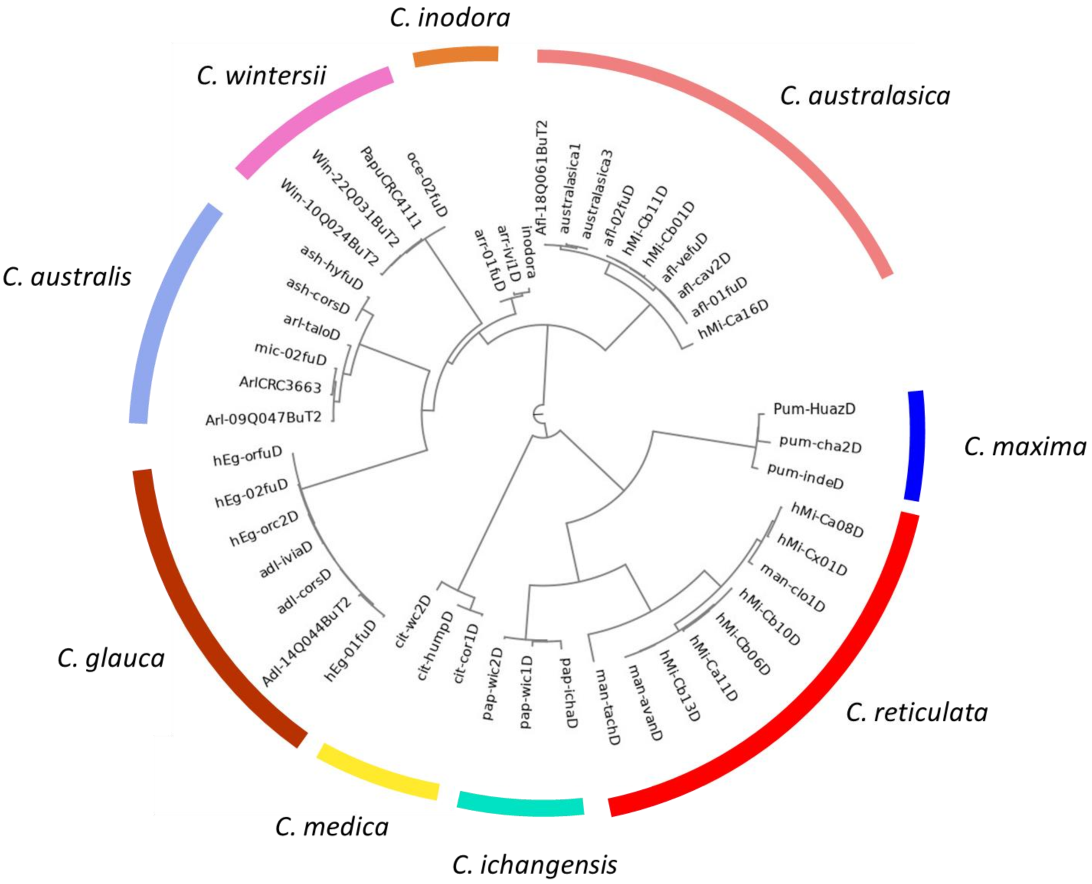


**Supplementary Figure 12:** Chloroplast inheritance of the analysed Oceanian admixed accessions

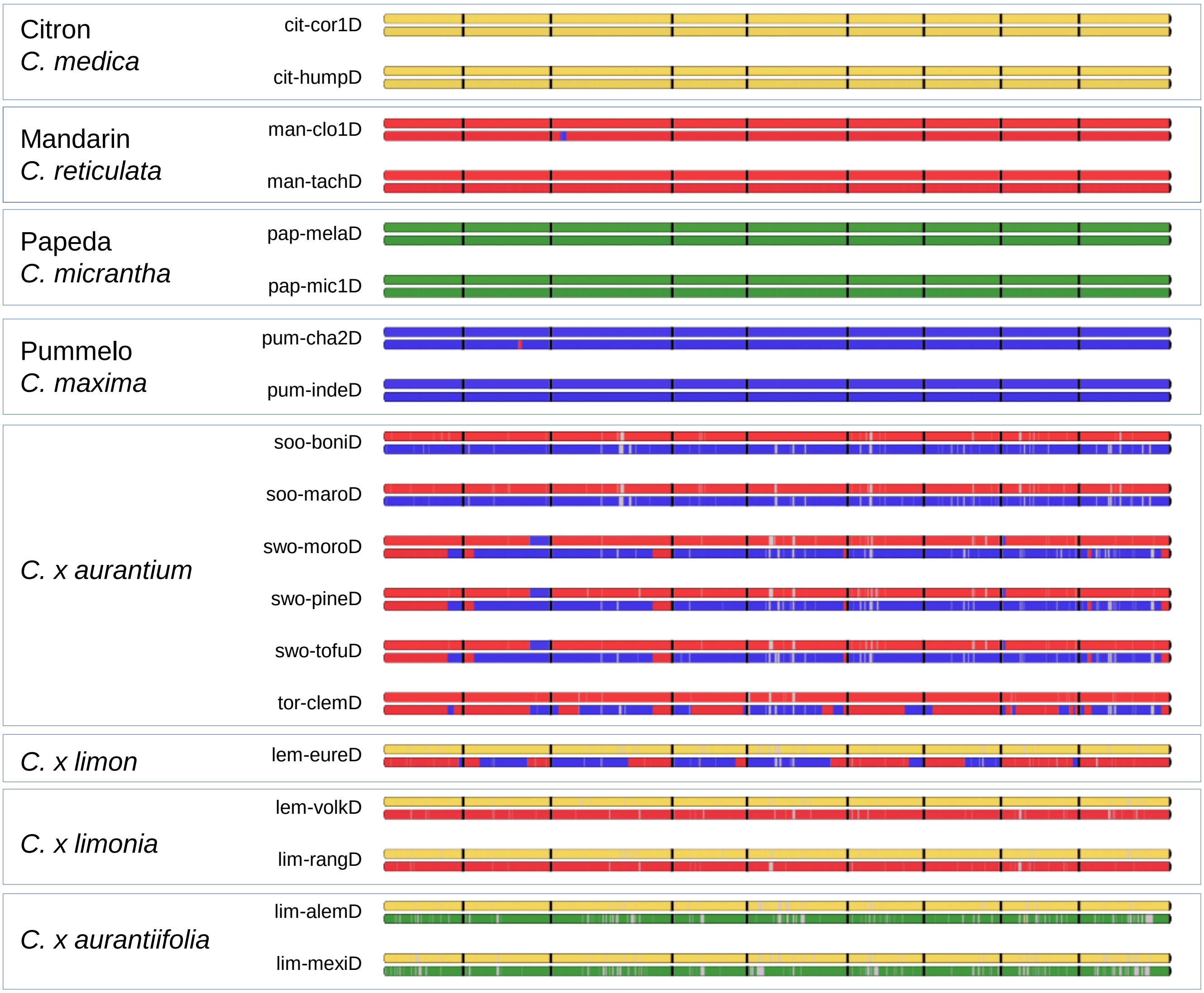


**Supplementary Figure 13:** Examples of Phylogenomic mosaic structure along the nine chromosome (separated by black dash) of some accessions representatives of the main Asian clades and admixed horticultural groups.

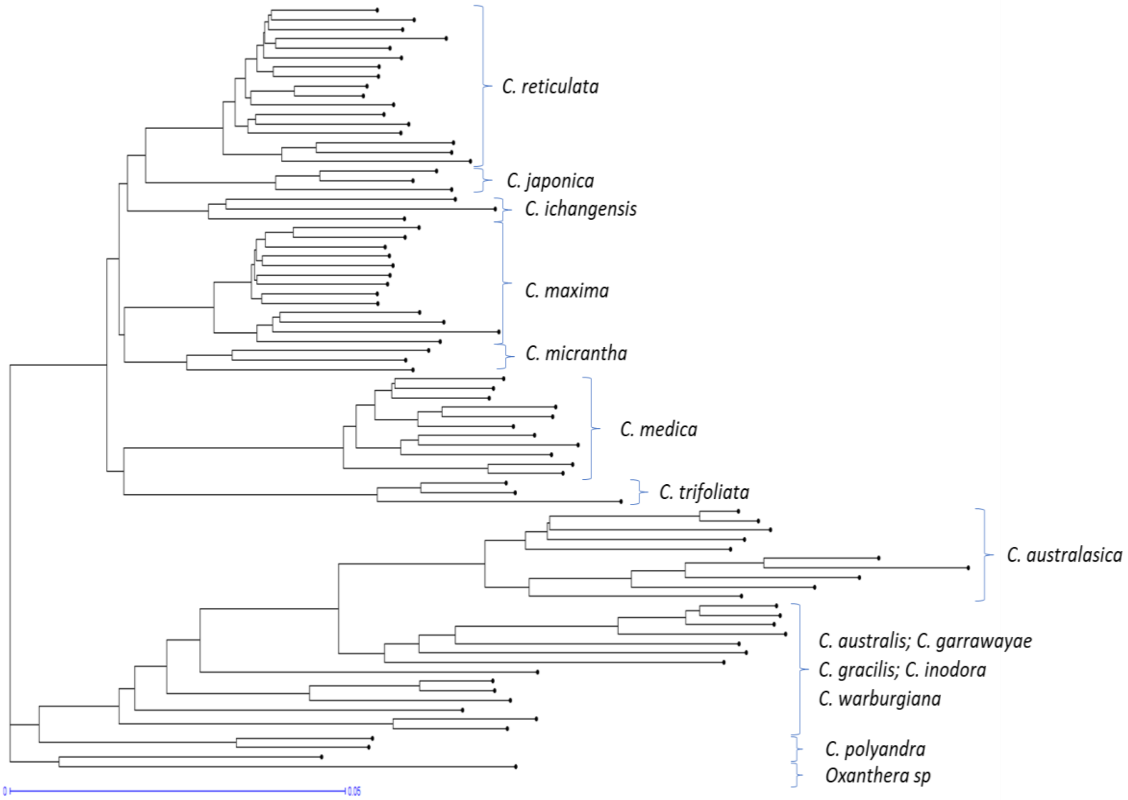


**Supplementary Figure 14. Neighbor-Joining tree based on gene presence/absence variation (PAV) across 81 citrus accessions.** The tree was constructed from binary gene loss data using SGSGeneLoss outputs mapped to the *AFL SRA 1002* reference genome.

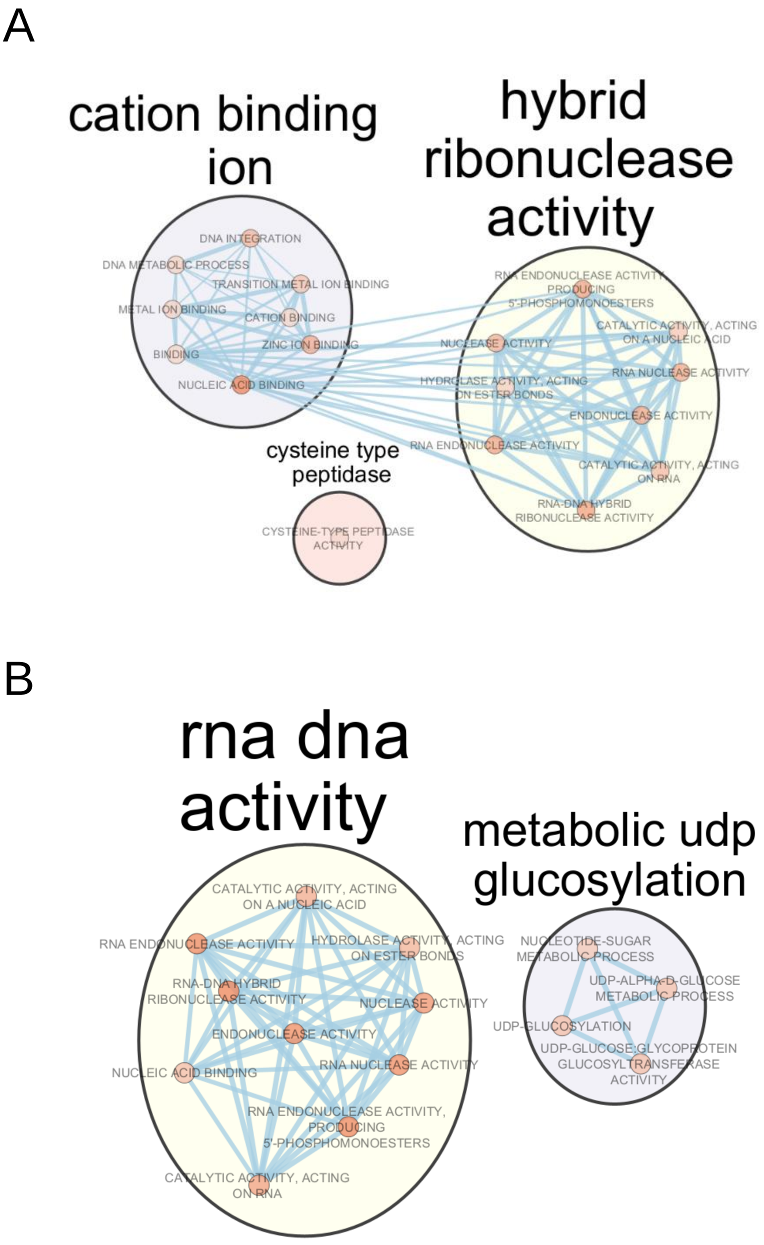


**Supplementary Figure 15. Gene Ontology (GO) enrichment networks of genes absent in Asian Citrus and conserved in Oceanian lineages. A)** GO enrichment network of the 127 genes present in ≥95% of Oceanian Citrus accessions but absent in all Asian species. **B)** GO enrichment network of the 52 genes exclusive to *C. australasica*.
